# Condensate screening identifies YM155 as β-catenin condensate inhibitor in colorectal cancer

**DOI:** 10.1101/2025.01.13.632724

**Authors:** Caterina Manzato, Nafiseh Sirati, Bronwen A. Knol, Hendrik J. Kuiken, Ben Morris, Cassio Fleming, Roderick L. Beijersbergen, Jurian Schuijers

**Affiliations:** Center for Molecular Medicine, University Medical Center Utrecht, Utrecht, The Netherlands; Oncode Institute, Utrecht, The Netherlands; Division of Molecular Carcinogenesis, Robotics and Screening Center, The Netherlands Cancer Institute, Amsterdam, The Netherlands

## Abstract

β-catenin is a transcriptional cofactor crucial in forming biomolecular condensates that drive WNT target gene transcription. Its pathological accumulation in the nucleus is a critical step in WNT-driven cancers, such as colorectal cancer. Disrupting β-catenin condensates is a promising cancer treatment strategy, but no clinical applications currently exist. In this study, we developed a Nanobody Enabled Condensate Observation (NECO) screening platform to identify small molecules that modulate β-catenin condensates. The platform revealed compounds that either enhanced or reduced β-catenin transcriptional condensates. Among them, YM155 (sepantronium bromide) was identified as an inhibitor, disrupting essential weak interactions for condensate formation. This disruption effectively inhibits the proliferation of WNT-driven colorectal cancer organoids. These findings show that the NECO platform can identify small molecules that target β-catenin condensates and suggest YM155 as a potential anti-cancer agent for colorectal cancer.

## INTRODUCTION

Aberrant activation of the WNT signaling-pathway^1,2^ is observed in more than 90% of colorectal cancers^3^ and is considered one of the key drivers of this cancer. The WNT-signaling pathway revolves around the control of the transcriptional activator β-catenin. In the absence of a WNT-ligand, β-catenin is continuously degraded but when the pathway is activated this degradation is halted and the accumulated β-catenin translocates into the nucleus where it activates transcription^4,5^. The WNT-pathway is a highly complex pathway with many homologous ligands, receptors, internal signal transducers, and DNA-bound transcription factors capable of compensating for chemical inhibition of other pathway components. However, this highly complex signaling network converges on β-catenin as the single transcriptional co-activator, making it a highly promising drug target. Despite its potential as a drug target no effective β-catenin inhibitors have been developed to date^6^, in part because β-catenin lacks easily druggable pockets.

To facilitate gene expression, the transcriptional machinery is locally concentrated at active genes in biomolecular condensates^7,8^. These condensates contain RNA-polymerase II, Mediator complex and transcription factors^9–11^ and we have shown that β-catenin also partitions into these condensates to activate WNT-driven transcription^12,13^. Condensate formation by β-catenin thus provides an opportunity to inhibit WNT-driven transcription that has not been explored.

Here we use our previously developed Nanobody Enabled Condensate Observation (NECO) platform^13^ to monitor β-catenin condensates in colorectal cancer cells and screen a library of small molecules. We find YM155 (sepantronium bromide) as a potent inhibitor of β-catenin condensation and WNT-driven transcription. We identify β-catenin as the molecular target of YM155 and show that YM155-mediated WNT inhibition is sufficient to selectively halt proliferation of patient derived colon cancer organoids. These findings show that transcriptional condensates can be targeted by small molecules to inhibit oncogenic transcription and identify YM155 as a β-catenin condensate inhibitor.

## RESULTS

### Screening of a chromatin targeting library using NECO platform identifies β-catenin condensate-modifying compounds

We have previously found that disruption of β-catenin condensates by co-factor derived peptides inhibited WNT-driven transcription^13^. To find small molecules that may have the same effect, we employed the NECO-β-catenin platform in HCT116 colorectal cells (Figure 1A) and tested a library of 216 compounds targeting proteins involved in chromatin maintenance, regulation, and gene expression. We limited the treatment time to 1 hour to select for compounds that can directly modify β-catenin condensates and minimize confounding secondary effects. After treatment, cells were fixed, and confocal images were acquired (Figure 1B). Custom image analysis allowed us to determine the abundance and size of the β-catenin condensates, as well as GFP signal intensity as a proxy for β-catenin concentration. This yielded both enhancers and reducers of nuclear β-catenin condensates (Figure 1C and Figure S1A). Compounds were selected for follow up based on their ability to modify condensate abundance, measured as condensates-per-nucleus z-score, and condensate intensity, measured as integrated intensity. They were further selected based on the reproducibility between replicate assays, measured as the standard deviation between the condensates-per-nucleus z-scores. Drugs with excessive variability and with disparities between condensates-per-nucleus and integrated intensity were excluded. We selected compounds that scored at the top of the distribution in terms of Mahalanobis distance (Figure S1B), meaning they scored as some of the most distant values from the center of the distribution generated combining z-scores from integrated intensity and condensates-per-nucleus.

**Figure 1:**
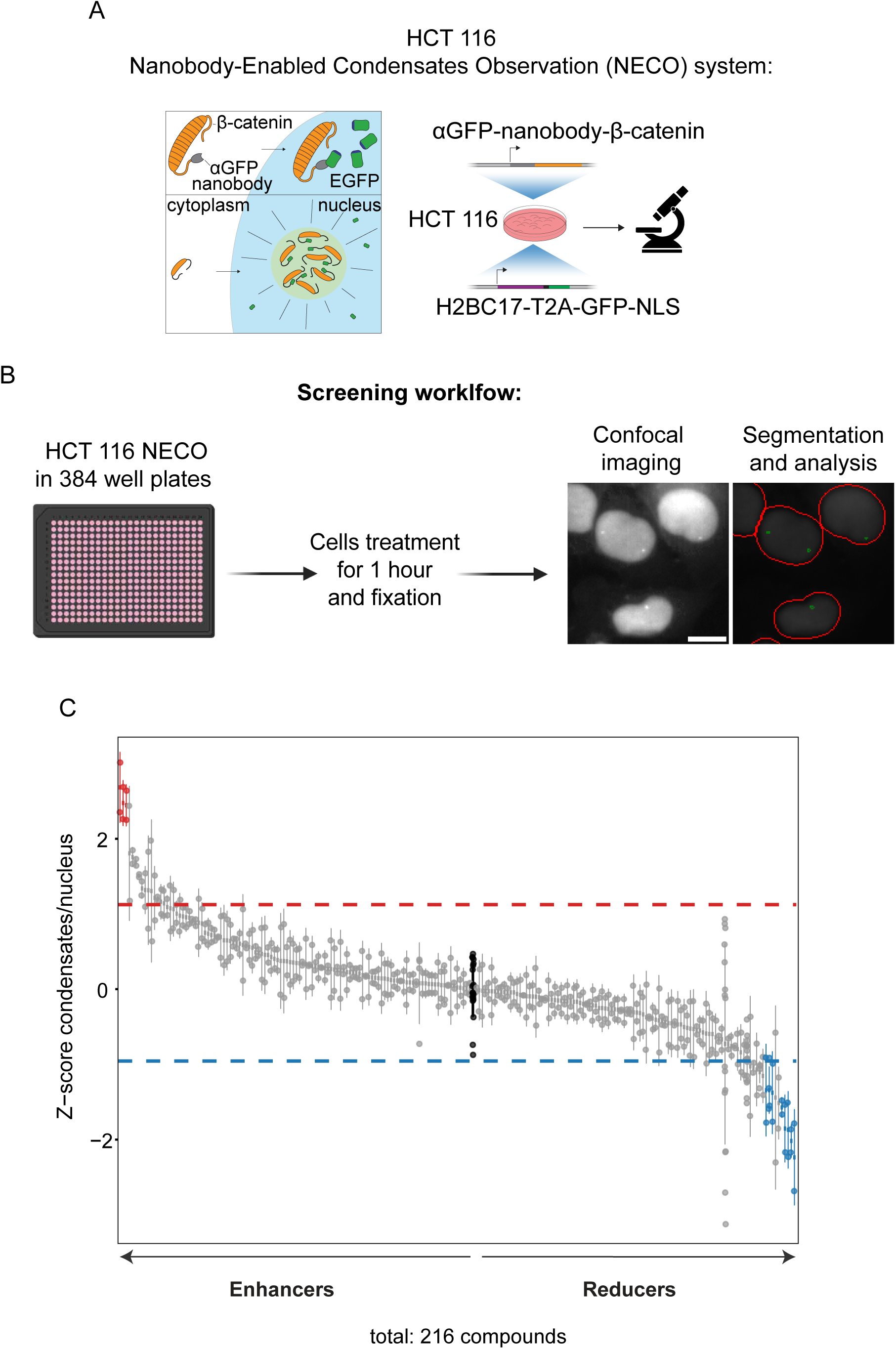
The Nanobody-Enabled Condensates Observation (NECO) identifies small molecule compounds targeting β-catenin condensates. A) Schematic representation of the Nanobody Enabled Condensates Observation (NECO) platform for β-catenin in HCT 116 cells. B) Schematic representation of the screening workflow C) Quantification of the number of condensates per nucleus z-score for all the 216 screened compounds and the DMSO control (black). Dotted red and blue lines show the limits of the 90% interval containing the untreated data (not shown). Enhancers hits are showed in red while reducers hits are shown in blue. Each point represents a biological replicate, at least two biological replicates were considered for each condition, 9 images were acquired for each biological replicate.

This process identified eight reducers and three enhancers, that qualified all our criteria. These data show that the NECO platform can identify compounds that modulate β-catenin condensates in the nucleus of colorectal cancer cells.

### Specific β-catenin condensate modulation by select compounds

To validate the effect on β-catenin condensates of the identified enhancers and reducers we performed titrations at concentrations 1 µM, 2.5 µM and 5µM, using the same NECO platform and treating for either 30 or 60 minutes. We quantified the average number of condensates per nucleus and the average condensed fraction per nucleus, measured as the sum of the integrated intensity of the condensates (Figures 2A, 2B and S2A). Based on these analyses, we identified a total of six compounds having a statistically significant effect on β-catenin condensates. Typically, compounds showed the strongest effects after 60 minutes treatment time, while still showing an appreciable effect after 30 minutes, with exception of BIX01294. Through these experiments we identified 3 enhancers: CYC116, CCT129202 and Pacritinib, and 3 reducers: YM155 (sepantronium bromide), WP1066, and BIX01294 (Figures 2A, 2B and S2B). Both CYC116 and CCT129202 are reported as Aurora Kinases inhibitors^14^ while both Pacritinib and WP1066 are known to target JAK2^15,16^. YM155 is reported as an inhibitor of survivin (*BIRC5*) transcription^17^ but its molecular target and the mechanism through which it regulates *BIRC5* expression is unclear. Finally, BIX01294 is reported to inhibit G9a histone methyltransferase^18^. All selected compounds are small aromatic molecules (Figure S2B). Feature analysis of the hits molecular descriptors using the rcdk package^19^ revealed no striking differences between the reducers and enhancers, except a slight increased hydrophilicity of YM155 compared to the other compounds (Supplementary Table 1). Hydrophilicity was measured by different computational prediction of the octanol-water partition coefficient (logP).

**Figure 2:**
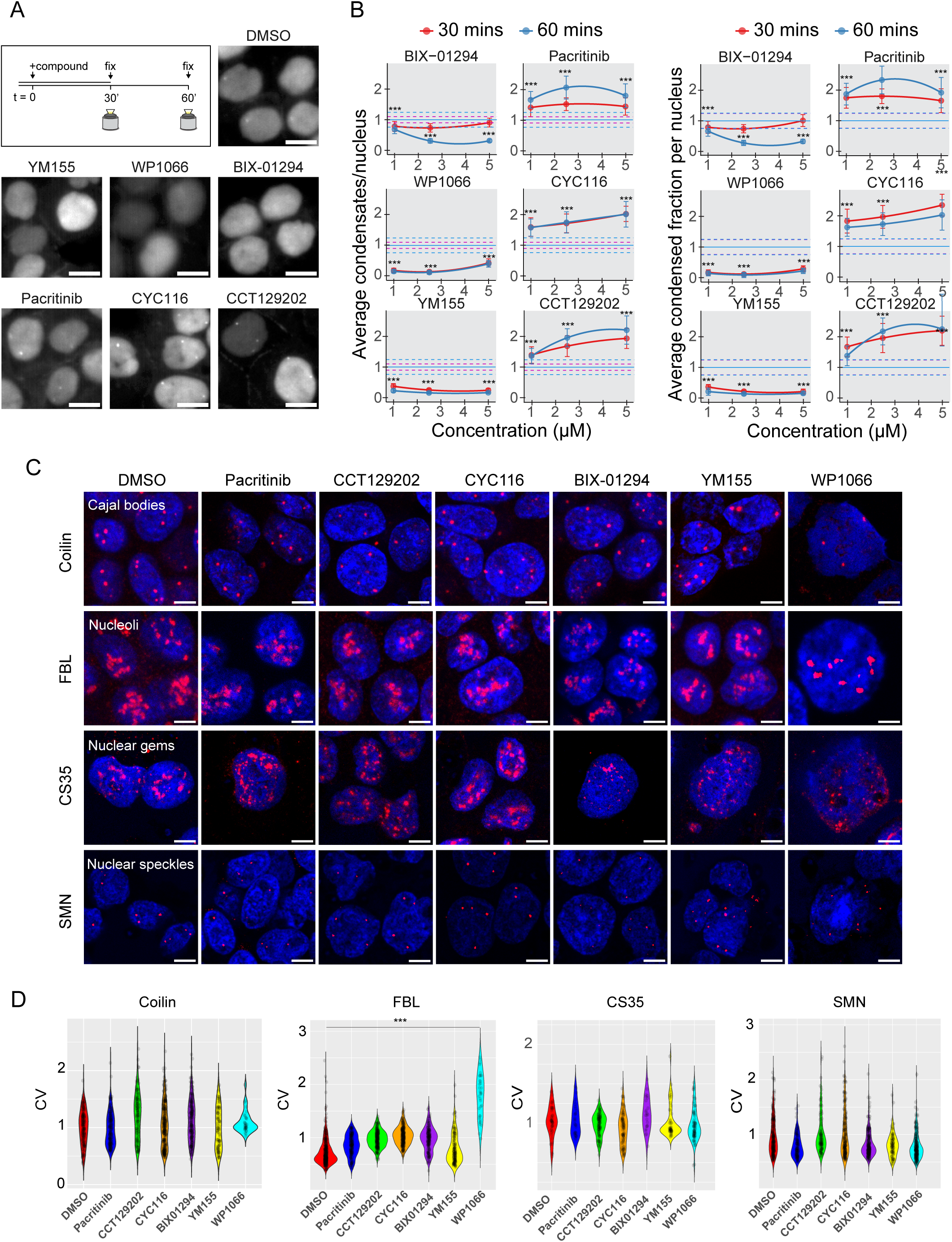
Titration experiment and staining for different condensates identify compounds that specifically affect β-catenin condensates. A) Schematic of the experiment and representative images of β-catenin condensates in the nuclei of HCT 116 NECO cells after 1 hour treatment with 0,01% DMSO and YM155, WP1066, BIX-01294, Pacritinib, CYC116 and CCT129202 (5 μM). B) Left: Quantification of the average number of β-catenin condensates per nucleus of HCT 116 NECO cells after 30 minutes (red) and 1 hour (blue) treatment with 0,01% DMSO and YM155, WP1066, BIX-01294, Pacritinib, CYC116 and CCT129202 (1 μM, 2.5 μM and 5 μM). Right: Quantification of the average condensed fraction of β-catenin condensates per nucleus in the of HCT 116 NECO cells after 30 minutes (red) and 1 hour (blue) treatment with 0,01% DMSO and YM155, WP1066, BIX-01294, Pacritinib, CYC116 and CCT129202 (1 μM, 2.5 μM and 5 μM). Continuous lines at y=1 represent DMSO controls, while dotted red and blue lines represent the error bars interval for DMSO at 30 minutes and 1 hour, respectively. At least five biological replicates for each treatment for each concentration were considered. 9 images were acquired for each biological replicate. One-way analysis of variance (ANOVA) and Tukey’s Honest Significant Difference (HSD) post hoc test were performed for each concentration. *p <0.05, **p <0.01, ***p < 0.001. C) Representative images of the immunofluorescence experiments for Colilin, FBL, CS35 and SMN on the HCT 116 NECO cells upon treatment with DMSO, YM155, WP1066, BIX-01294, Pacritinib, CYC116 and CCT129202. Scalebar: 5 μm. D) Quantification of the covariate of the immunofluorescence experiment for Colilin, FBL, CS35 and SMN on the HCT 116 NECO cells upon treatment with DMSO, YM155, WP1066, BIX-01294, Pacritinib, CYC116 and CCT129202. The coefficient of variation of each image and subsequently the mean CV (std Intensity/Mean Intensity) of each condition were calculated and plotted using R. A one-way ANOVA was conducted to assess the impact of drug treatment. Significant differences between drug treatments were further explored using Tukey’s HSD post hoc test. ***p < 0.001. A total of 10 images for each treatment were collected from 2 biological replicates from two independent experiments.

To determine the specificity of the selected of compounds, we tested whether they affect other well-defined nuclear condensates. Using immunofluorescence to monitor Nucleoli (FBL), Cajal bodies (Coilin), Nuclear gems (CS35) and Nuclear speckles (SMN) we found that none of the compounds have an effect on the tested nuclear condensates, except WP1066 that affected nucleoli and the shape and size of the treated cells. (Figures 2C, 2D). This may be explained by rapid cytotoxic effects of WP1066 at the tested concentrations and indicated that the observed effects on β-catenin condensates are likely not specific for this compound. Together, these results suggest CYC116, CCT129202, Pacritinib, YM155 and BIX01294 specifically modify β-catenin condensates in HCT116 colorectal cancer cells.

### YM155, Pacritinib and CCT129202 directly affect β-catenin homotypic droplet formation

Next, we asked whether the selected compounds could influence β-catenin homotypic interactions directly. To do so, we tested the compounds in an *in vitro* droplet assay with purified recombinant mEGFP-β-catenin^13^. Condensed fraction and partition ratio quantifications showed reducer YM155, and enhancers Pacritinib and CCT129202 to also reduce and enhance homotypic β-catenin droplets *in vitro* (Figure 3A and 3B) at stoichiometry 3:10 mEGFP-β-catenin:compound. Taken together, these experiments show that YM155, Pacritinib and CCT129202 can affect β-catenin homotypic droplet formation *in vitro*.

**Figure 3:**
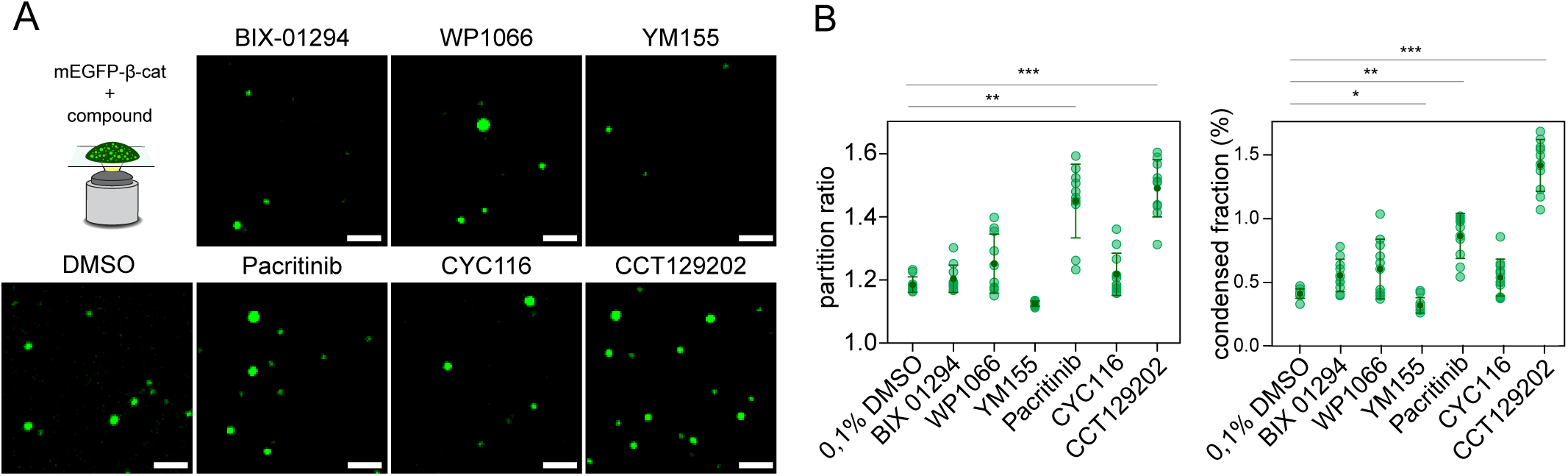
YM155, Pacritinib and CCT129202 affect β-catenin homotypic interactions *in vitro*. A) Schematic representation and representative images of droplet formation assays of 3 μM purified mEGFP-β-catenin with 10 μM of the compounds: YM155, WP1066, BIX-01294, Pacritinib, CYC116 and CCT129202. 0,1% DMSO was used as a control. Scalebar: 2 μm. B) Quantification of the droplet formation assay of 3 μM purified mEGFP-β-catenin with 10 μM of the compounds: YM155, WP1066, BIX-01294, Pacritinib, CYC116 and CCT129202. 0,1% DMSO was used as a control. 10 images for each condition were analysed. Kruskal-Wallis test and Dunn’s multiple comparison test were performed to assess the effect of the small molecule on β-catenin droplets partition ratio and condensed fraction. *p <0.05, **p <0.01, ***p < 0.001.

### YM155 Inhibits β-catenin-driven transcription

Condensates are highly dynamic structures that can rapidly adapt to changing conditions. To investigate the prolonged effect of the selected compounds on the HCT 116 NECO cell line we performed a live-imaging time course experiment: acquiring images every 30 minutes starting 1 hour after the treatment, for 12 hours. All 6 compounds recapitulated the effects of our short 60-minute treatment, but condensate numbers changed considerably during prolonged exposure. Of the three reducers, YM155 performed the best, as it reduced β-catenin condensate numbers for around 5 hours after the 1 μM treatment, while the other two reducers showed a shorter condensate reduction before condensate numbers returned to control levels (Figure 4A, 4B and S4A). Among the enhancers, Pacritinib showed the highest increase of the condensates per nucleus with a sustained effect for 5 hours after treatment. CCT129202 and CYC116 showed a milder increase of the number of condensates per nucleus that was maintained for 2 and 3 hours, respectively (Figure 4A, 4B and S4A). Of note, the average number of nuclei per image was reduced by YM155, starting at 12 hours after the treatment, concomitantly with the increase in the number of condensates in the nuclei (Figures 4B and S4B). For this reason, we interpret the increased condensates number at that time point to be the result of decreased viability of the cells, rather than reduced compound concentrations. These data show that condensate modifying effects reduce in strength over time, with YM155 being the most potent condensate reducer out of the tested compounds.

**Figure 4:**
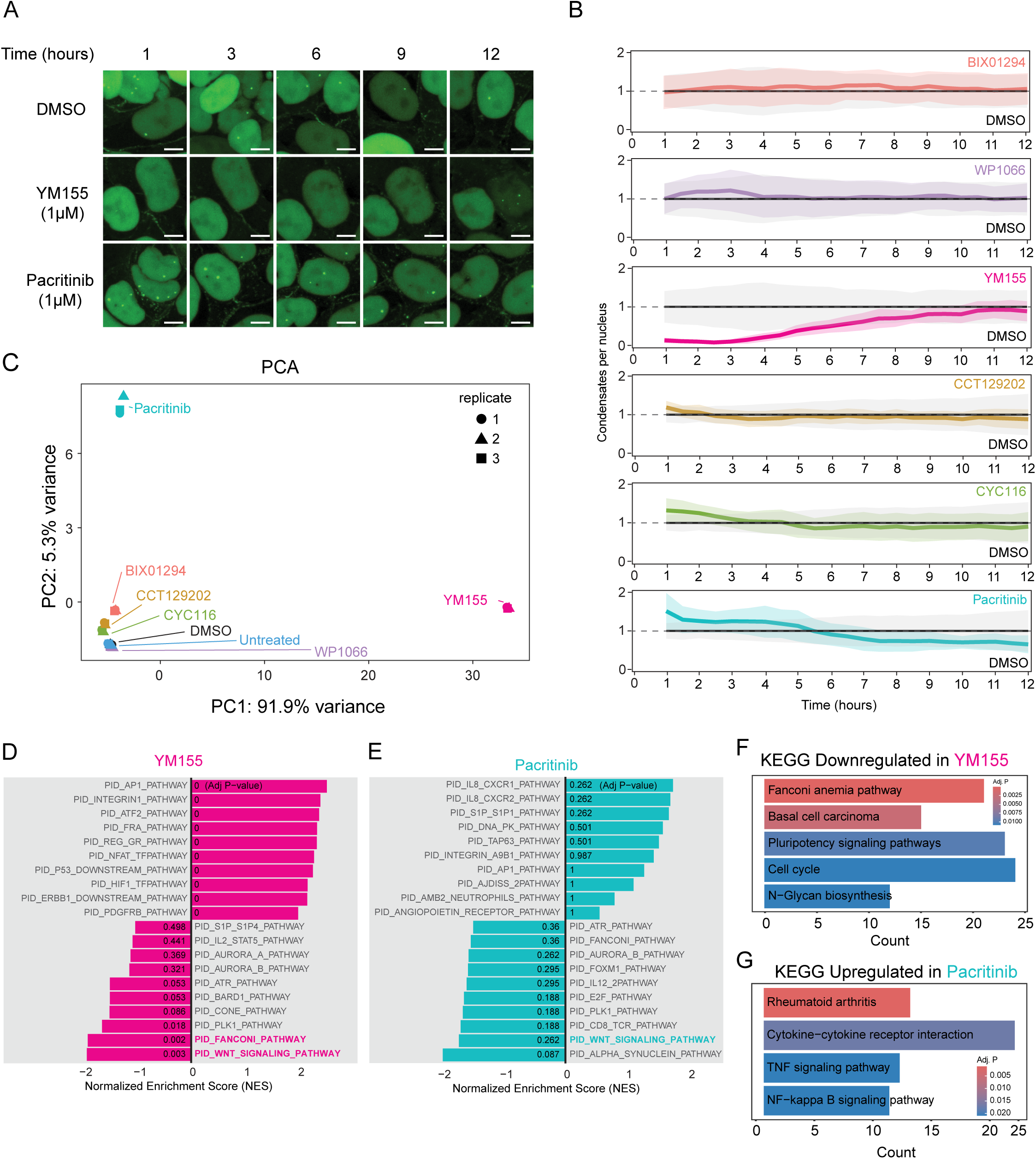
Live imaging timelapse and RNA-sequencing experiments link condensate dynamics and transcriptional output. A) Representative images of live HCT 116 NECO cells at 1 hour, 3 hours, 6 hours, 9 hours and 12 hours timepoints after 0,01% DMSO, 1μM YM155 and 1μM Pacritinib treatments. Scalebar 5μm. B) Quantification of the average number of condensates per nucleus over time upon the treatment with 0,01% DMSO and six compounds: YM155, WP1066, BIX-01294, Pacritinib, CYC116 and CCT129202 (1μM). A total of 16 images for treatment were collected from a total of 4 biological replicates from two independent experiments. C) Principal component analysis of the transcriptome analysis of RNA samples obtained after 12 hours of treatment with 0,01% DMSO and 1μM YM155, WP1066, BIX-01294, Pacritinib, CYC116 and CCT129202. Treatments were performed in triplicates. D) Gene set enrichment analysis for HCT 116 cells transcriptome upon 12 hours 1μM YM155. Adjusted p-value indicated inside the plot bars. Treatments were performed in triplicates. E) Gene set enrichment analysis for HCT 116 cells transcriptome upon 12 hours 1μM Pacritinib. Adjusted p-value indicated inside the plot bars. Treatments were performed in triplicates. F) Enriched pathways from KEGG database pathways analysis on the significatively downregulated genes upon 12 hours 1μM YM155 treatment. Treatments were performed in triplicates. G) Enriched pathways from KEGG database pathways analysis on the significatively downregulated genes upon 12 hours 1μM Pacritinib treatment. Treatments were performed in triplicates.

In a previous study, we demonstrated that β-catenin condensate dissolution results in reduced transcription of WNT target genes^13^. To determine if the modulation of β-catenin condensates by these compounds also resulted in WNT target gene changes, we performed transcriptome analysis 12 hours after applying a 1μM treatment. In agreement with the live-imaging results, Pacritinib and YM155 had stronger effects than the other compounds, their profiles clustering separately from all the other samples in a principal component analysis (Figure 4C). To better understand the upregulated and downregulated genes affected by these two treatments we performed gene set enrichment analysis. Gene set enrichment analysis using Panther^20^ gene sets found the WNT signaling pathway significantly enriched among the YM155-downregulated genes (Figure 4D). Unexpectedly, the same Panther WNT-pathway was also reported among the downregulated pathways for Pacritinib, although the associated p-adjusted value is above the 0.05 cutoff (0.262) (Figure 4E). We reason that, while Pacritinib enhanced β-catenin condensate numbers, these condensates may be altered functionally, leading to the downregulation of WNT target genes. These results indicate that both reducer YM155 and enhancer Pacritinib are inhibiting WNT-target gene expression (Figure S4C), in spite of opposite effects on β-catenin condensate numbers.

To further investigate the transcriptional changes induced by these compounds we performed KEGG^21^ pathway analysis for significant downregulated genes in YM155, which showed a strong enrichment of genes involved in the Fanconi Anemia pathway (Figure 4F). The effect of YM155 on Fanconi Anemia pathway genes was previously reported^22^, although the mechanism through which this happens is unclear. Besides Fanconi Anemia, “cell cycle regulation” and “signaling pathways regulating stem cell pluripotency” were enriched, which are in line with the perturbation of WNT signaling^23,24^ (Figure 4F). Other databases (Reactome^25^ and Gene Ontology^26,27^) used for pathway analyses also supported the link between YM155 and Fanconi Anemia pathway, cell cycle genes, and WNT pathway-regulated processes (Figure S4D and S4E). The same analyses of the Pacritinib-induced downregulated genes, using KEGG and Reactome databases, did not show any significantly enriched pathway, while the Gene Ontology pathway analysis showed two significantly downregulated pathways being: axon guidance and neuron projection guidance both with a padj of 0,014. Conversely, Pacritinib-upregulated genes were enriched for “regulation of the immune response”, which is in accordance with the reported events downstream the inhibition of STAT3 in cancer^28^ (Figure 4G). These analyses show that YM155 downregulates genes involved in the Fanconi Anemia pathway, as well as the WNT-pathway. The two pathways have been linked in breast cancer, where WNT-pathway inhibitors can down-regulate DNA repair associated genes and create a BRCA-like state^29^. These data also identify WNT pathway regulation patterns in Pacritinib treated samples, although they are masked by the inhibition of the JAK2/STAT3 downstream programs.

### YM155 induces cell death in colorectal cancer cells in a β-catenin dependent manner

We have previously found that perturbation of β-catenin condensates inhibits proliferation of HCT116 colorectal cancer cells and here we observed an effect of YM155 on cell viability in our live imaging experiment (Figure S4B). We aimed to investigate the effect of YM155-induced β-catenin condensate perturbation on the induction of cell death. β-catenin harbors two Intrinsically Disordered Regions (IDRs), one located at the C-terminal and one at the N-terminal, that have been shown to be fundamental for the formation of condensates^12,13^. Purified recombinant GFP-β-catenin^Wt^ is forming droplets in vitro, while GFP-β-catenin^IDRs*^, GFP-β-catenin^cIDR*^ and GFP-β-catenin^nIDR*^ are less capable of forming droplets in vitro than the wildtype^30^. To investigate importance of β-catenin condensation, we employed HCT116 β-catenin KO and derived cell lines re-expressing condensate forming (mSc-β-catenin^Wt^) and condensate-attenuated forms of β-catenin (mSc-β-catenin^nIDR*^, mSc-β-catenin^IDRs*^ and mSc-β-catenin^cIDR*^). We studied the effect of YM155 on these cell lines in a dose-dependent manner where samples were subjected to propidium iodide exclusion assay 24 hours after the treament. Results of dose-response curves show a higher cell death in the β-catenin^Wt^ cell line with respect to the other four cell lines considered (figure 5A), suggesting a role for β-catenin condensates in the induction of YM155-induced cell death. Of note, the mSc-β-catenin^nIDR*^ showed an intermediate level of cell death between the β-catenin^Wt^ line and the other lines. YM155 was able to induce cell death in these cell lines at concentrations as low as 4nM, 8nM and 16 nM YM155. To test the effect on condensates of nanomolar concentrated YM155, HCT 116 NECO cells were imaged 24 hours following treatment with 4nM, 8nM and 16nM YM155 (Figures 5B and 5C). Results revealed a significant reduction of condensates abundance compared to the DMSO control.

**Figure 5:**
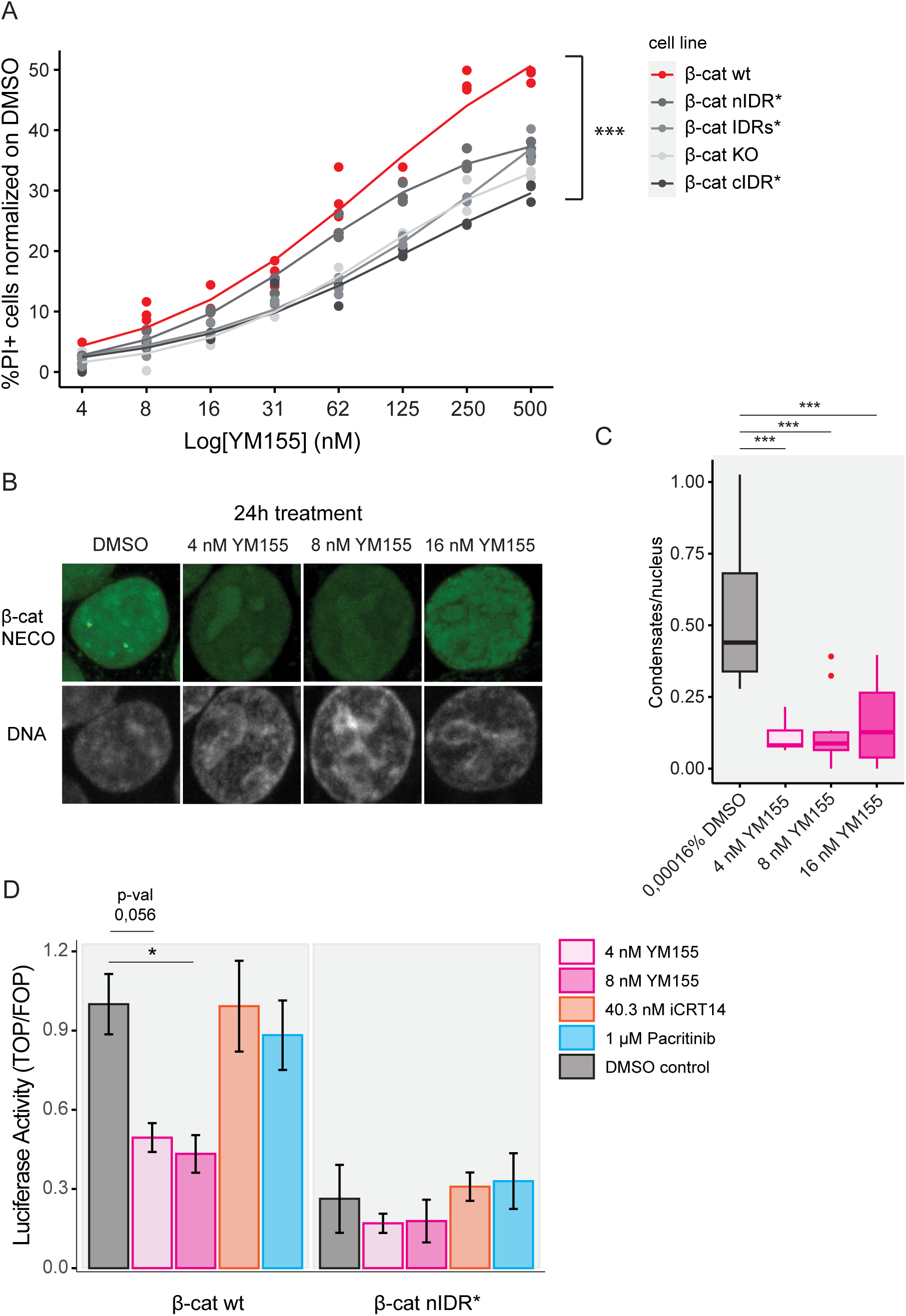
YM155 works as β-catenin condensate reducer and WNT transcriptional modulator in the nanomolar range. A) Quantification of YM155-induced death at different YM155 concentrations and on different cell lines: mSc-β-catenin^Wt^ (β-cat^Wt^), mSc-β-catenin^nIDR*^ (β-cat^nIDR*^), mSc-β-catenin^IDRs*^ (β-cat^IDRs*^), β-catenin KO (β-cat KO), mSc-β-catenin^cIDR*^ (β-cat^cIDR*^). Cells were treated for 24 hours. Percentage of propidium iodide (PI)-positive cells is normalized on DMSO. Experiments were performed in four independent experiments (two concentrations for each experiment) in triplicates. One-way ANOVA and Tukey’s HSD post hoc test on the upper limit of the curves (500 nM) were performed. Comparisons between β-cat^Wt^ and the other three cell lines were ***. ***p < 0.001. B) Representative images of the effect of 24 hours treatment on β-catenin condensates in HCT 116 NECO cells with 4 nM, 8 nM and 16 nM YM155. DMSO 0,00016% was used as a control. C) Quantification of the effect of 24 hours treatment on β-catenin condensates in HCT 116 NECO cells with 4 nM, 8 nM and 16 nM YM155. DMSO 0,00016% was used as a control. Welch’s ANOVA and Games-Howell post hoc test were performed. A total of 10 images for condition from a total of 2 replicates were used for the analysis. ***p < 0.001. D) Luciferase assay on mSc-β-catenin^Wt^ (β-cat^Wt^), mSc-β-catenin^nIDR*^ (β-cat^nIDR*^) treated with 4 nM YM155, 8 nM YM155, 40.3 nM iCRT14, 1 μM Pacritinib and their respective DMSO controls. All the samples were normalized on their DMSO controls. 0,00004% DMSO is showed. Data is normalized to the mSc-β-catenin^Wt^ (β-cat^Wt^) DMSO control. Two-way ANOVA and Tukey’s HSD post hoc tests were performed. A total of six replicates from 2 independent experiments were considered. *p <0.05.

To test if YM155 affects β-catenin-driven transcription in a condensate dependent manner we assessed the transcriptional output upon YM155 treatment using the mSc-β-catenin^Wt^ and mSc-β-catenin^nIDR^ cell lines, as mSc-β-catenin^nIDR^ is the only mutant line showing residual WNT transcriptional activity (Figure S5A). Cells were treated with 4nM and 8nM YM155, 1 µM Pacritinib, and the β-catenin inhibitor iCRT14 at the reported IC50 of 40.3nM^31^. We found that YM155 reduced β-catenin transcriptional activation by approximately 50% in mSc-β-catenin^Wt^ cells, while no significant reduction was observed in the mSc-β-catenin^nIDR^ expressing line. Interestingly, the reported β-catenin inhibitor iCRT14 did not show an effect at tenfold higher concentration than YM155. Pacritinib treatment did not show any significant effect, in line with the previous RNA-seq results. Taken together, these results indicate the relevance of β-catenin condensates in the induction of cell death upon YM155 treatment. Furthermore, we observed a correlation between cell death, β-catenin condensate reduction, and impaired WNT specific transcription, when β-catenin proficient cells were treated with nanomolar concentrated YM155.

### YM155 selectively affects viability of APC mutated colon organoids

Due to the important role of the WNT-pathway in tissue maintenance, many compounds targeting the WNT-pathway exhibit significant off-target toxicity. However, WNT-driven colorectal cancer has elevated levels of β-catenin specifically in cancer cells that may differentiate between healthy and diseased cells^32^. To assess the potential of YM155 as an anticancer therapeutic, we tested the drug on isogenic healthy and APC mutant colon organoids. APC plays a fundamental role in the WNT pathway as part of the destruction complex^33^. Its mutation leads to a non-functional destruction complex and to the accumulation of β-catenin in the nucleus. We treated both organoid lines with increasing concentrations of YM155 and we measured the cell death with a dual fluorescent dye assay (Figures 6A and 6B). We observed that organoids bearing the APC mutation are display more cell death marker, induced by YM155, indicating a potential therapeutic window in which YM155 preferentially affects APC transformed colon cancer cells. Altogether, these data show that b-catenin condensate formation might be preferentially perturbed in APC-mutated cancer cells by YM155.

**Figure 6:**
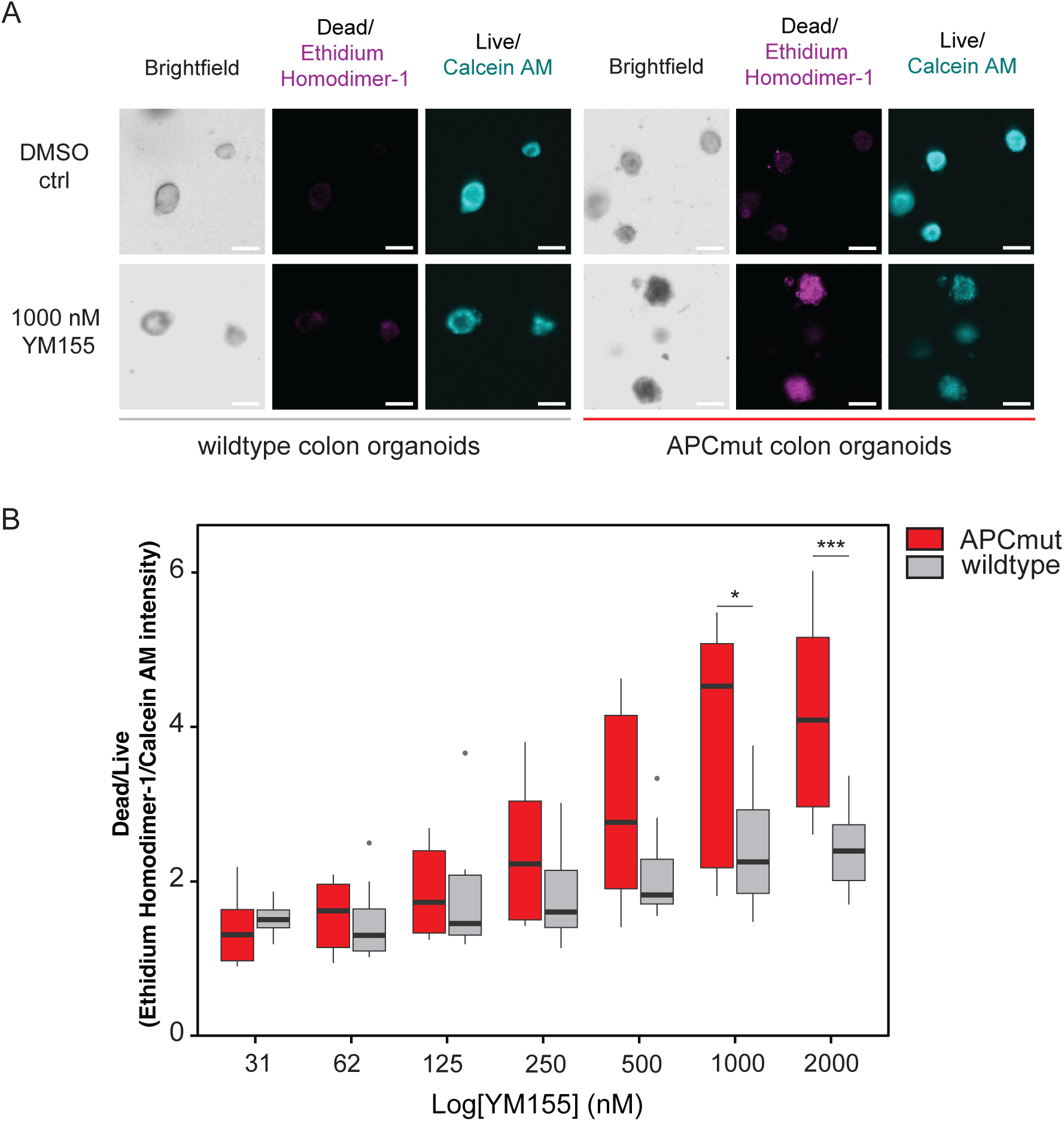
YM155 affects viability of colon cancer organoids to a higher extent than wildtype colon organoids. A) Example images of live/dead staining with the Ethidium Homodimer-1/Calcein AM method on colon organoids. Wildtype and APC mutant organoids are treated with 0,01% DMSO or 1 μM YM155 for 24 hours. Scalebar: 100 μm. B) Quantification of the Ethidium Homodimer-1/Calcein AM ratio on colon organoids wildtype and APC mutated. Organoids were treated with the indicated concentrations of YM155 and their respective controls for 24 hours. 18 three channels-images for each condition were acquired from 6 biological replicates from 2 independent experiments for each genotype. Two-way ANOVA and Tukey’s HSD post hoc tests were performed. A total of six replicates from 2 independent experiments were considered. *p <0.05, ***p < 0.001.

## DISCUSSION

In this study we aimed to identify small molecules able to modify β-catenin condensates. To do so, we established a high-throughput imaging screening platform and tested a library of 216 chromatin targeting compounds. The NECO (Nanobody-Enabled Condensates Observation) platform permitted us to study the direct effect of the compounds on β-catenin condensate prevalence, intensity, size and shape, which cannot be determined by reporter-based or genetic screenings. Through this method and further validations, we identified YM155 as an effective β-catenin condensate modifier among the hits. It significantly reduces the number of β-catenin condensates in colorectal cancer cells, and it directly disrupts homotypic β-catenin droplets *in vitro*. Furthermore, YM155 was found to downregulate the expression of the WNT target genes in our transcriptome analyses.

YM155-induced cell death on cell lines bearing different mutations of β-catenin, allowed us to understand the role of β-catenin condensates in the mechanism. Condensate-attenuated β-catenin mutants were less sensitive than the wildtype counterpart, and to some extent than the mSc-β-catenin^nIDR*^ mutant, to YM155 induced cell death. Furthermore, the β-catenin mutant that showed some residual WNT transcriptional activity was less susceptible to the transcriptional inhibition by YM155 treatment than wild type β-catenin. Finally, YM155 was able to preferentially affect viability of human APC-mutated colon cancer organoids over wildtype colon organoids. This indicates the drug exhibits some specificity towards WNT-activated colon cancer, indicating YM155 may be a candidate for application in this cancer.

Taken together, our results indicate that β-catenin condensates represent a target of YM155. This compound was previously shown affecting the expression of the survivin gene^17^, although the mechanism through which it exerts this function was never reported. Of note, the concentration at which we find YM155 affecting viability of colorectal cancer cells HCT116 and organoids is comparable to the effective tested concentrations in a previous study^34^. This suggests survivin inhibition might happen downstream β-catenin inhibition by YM155. Moreover, we showed an effect of YM155 on Fanconi anemia pathway genes, previously showed also in the breast cancer cell line MDA-MB-231^22^. The same genes were found downregulated upon WNT inhibition in this study^29^, in line with our findings that YM155 acts as a WNT pathway inhibitor. However, the molecular details of the process through which YM155 binds to β-catenin, affects the condensate intermolecular weak interactions and thus leads to cell death, will require more study.

In conclusion, employing a screening for β-catenin condensate modifying compounds we identified a small molecule that directly binds to β-catenin, disrupts β-catenin condensates, inhibits β-catenin-driven transcription and preferentially induces apoptosis in APC-mutated colon cancer organoids. Although the mechanism of condensate inhibition needs to be further elucidated, these findings open new avenues for the identification of molecules that disrupt condensates and could provide some insights for the design of new compounds. Importantly, as clinical safety of YM155 was previously established^35^, this drug may be suitable for repurposing on colorectal cancer and other WNT-high cancers. Furthermore, we establish the NECO platform as a method that can be used for the identification of condensate modifiers for other types of nuclear condensates.

## MATERIALS AND METHODS

### Cell culture

The HCT116 NECO cell line was generated in the previous study^13^ where called HCT116 - H2BC17-T2A-EGFP GNb-b-catenin cell line. Cells were cultured in high-glucose DMEM (Sigma; D6429-24X500ML) supplemented with 10% FBS and Penicillin-Streptomycin at 37C and 5% CO_2_.

HCT 116 β-catenin KO, mSc-β-catenin^Wt^, mSc-β-catenin^IDRs*^, mSc-β-catenin^nIDR*^, mSc-β-catenin^cIDR*^ cell lines were generated as follows. Knock-out lines for β-catenin in HCT116 cells were generated by CRISPR-Cas9, by transfection of pSpCas9(BB)-2A-GFP (Addgene #48138) with gRNA sequence (5’-*GTTCCCACTCATACAGGACTTGG-3’)* targeting β-catenin, using Lipofectamine LTX. Two days following transfection, GFP-positive cells were sorted by FACS in 96-wells culture plates to obtain monoclonal lines. Clones were screened for successful knock-out of β-catenin by immunofluorescence and validated by both western blotting and sequencing of the genomic locus. Stable lines of HCT116 cells expressing indicated constructs were generated using lentiviral transductions followed by selection with Puromycin (Bio connect; SC-108071B) or Blasticidin (Bio Connect; ant-bl-1). In HCT116 β-catenin knockout lines in which mSc-β-catenin^Wt^ or mSc-β-catenin^IDRs*^ were re-expressed, cells were sorted by FACS to obtain monoclonal lines.

### Organoids culture

Organoids were cultured at 37°C and at 5% CO_2_. The basic culture medium contained advanced DMEM/F12 supplemented with 1% penicillin/streptomycin (Gibco), 10mM HEPES (Gibco), 1% Glutamax (Gibco), 1X B27 (Invitrogen), 50ng/ml Human recombinant EGF (Peprotech), 1,25 mM N-acetylcysteine (Sigma-Aldrich), 10mM Nicotinamide (Sigma-Aldrich), 500nM A83-01 (Tocris), 10µM SB202190 (Gentauer), 10% Noggin conditioned media. Wildtype colon organoids media was supplemented with 1,6nM Wnt surrogate Fc Fusion protein (Immuno Precise Antbodies) and 20% R spondin conditioned media. Organoids were splitted through Trypsin-EDTA (Sigma-Aldrich) treatment. Organoids were were cultured in drops of Basement Membrane Extract (BME; Amsbio) and cultured for the first two days after splitting with 10 μM Y-27632 dihydrochloride (Gentaur). The organoid line HUB-02 – A2 – 130 was used and the APC mutant was generated as previously reported^36^ (gRNA3) starting from this line.

### Screening

HCT 116 NECO cells were plated in 384 well plates in duplicate. 3500 cells for each well were plated 24 hours before the treatment. Cells were treated with a library of 216 compounds targeting the epigenome. Drugs used were diluted in DMSO at initial concentration of 10mM. 6% 1,6 hexanediol and 0,05% DMSO treated cells were used as controls. Treatment with the compounds was performed for 1 hour, with an exception for 6% 1,6 hexanediol treatment, that was performed for 5 minutes. After the treatment cells were fixed and stained in 3,7% PFA, 5 μg/ml Hoechst33342 in PBS and incubated for 15 minutes. Plates were washed 3 times with PBS. Cells were imaged with the automated confocal microscope ImageXpress micro 4 (Molecular Devices) with a 20X objective. 9 images for each well were acquired.

### Hits validation

HCT 116 NECO cells were plated in two 384 well plates 24 hours before the treatment. Treatments for 8 reducers: Tozasertib (VX-680) (SelleckChem), Danusertib (PH-739358) (SelleckChem), WP1066 (MedChem), YM155 (sepantronium bromide) (SelleckChem), Salermide (MedChem), BIX 01294 (SelleckChem), EPZ004777 (SelleckChem), MK-5108 (VX-689) (MedChem) and 3 enhancers: CYC116 (SelleckChem), CCT129202 (SelleckChem), Pacritinib (SB1518) (SelleckChem), were performed at 3 different concentrations (1 μM, 2.5 μM and 5 μM), for each concentration and treatment 5 replicate wells were treated. Drugs used were diluted in DMSO at initial concentration of 10mM. Tanshinone IIA (1 μM, 2.5 μM and 5 μM) and 0,05% DMSO were used as controls. Treatments were conducted for 1 hour in one of the plates and for 30 minutes in the other. Cells were fixed fixed and stained in 3,7% PFA, 5 μg/ml Hoechst33342 in PBS for 15 minutes. Subsequently, plates were washed 3 times in PBS. Cells were imaged with the automated confocal microscope ImageXpress micro 4 (Molecular Devices) with a 20X objective. 9 images for each well were acquired.

### Live imaging

8 x 10^4^ NECO cells were seeded in each well of a μ-slide 8 well chamber (ibidi) 24 hours before the experiment. Cells were treated with three enhancers CYC116 (SelleckChem), CCT129202 (SelleckChem) and Pacritinib (SB1518) (SelleckChem) or three reducers compounds WP1066 (MedChem), YM155 (sepantronium bromide) (SelleckChem) and BIX 01294 (SelleckChem). All drugs were used at 1 μM concentration and the 0,01% DMSO treatment was used as control. Cells were treated in duplicate wells and 4 images for each well were acquired with a 60X oil immersion objective on the Nikon spinning disc microscope. 29 stacks of 0.8 μm each were acquired for each image. Images were acquired every 30 minutes for 15 hours, starting 1 hour after the treatment.

### 24 hours YM155 treatment

8 x 10^4^ NECO cells were seeded in each well of a μ-slide 8 well chamber (ibidi) the day before the treatment. Cells were treated with 0,00016% DMSO, 16nM YM155, 8nM YM155, 4nM YM155. Cells were fixed with 2ug/ml Hoechst in 3,7% PFA in PBS for 15 mins and subsequently washed with PBS. Cells were treated in duplicate wells and 5 images for each well were acquired with a 60X oil immersion objective on the Nikon spinning disc microscope. 29 stacks of 0.8 μm each were acquired for each image.

### Image analysis

Images were analysed with a custom CellProfiler pipeline. Maximal intensity projections were created and then nuclei were segmented based on Hoechst33342 staining, images were masked for the nuclei segmentation and the condensates were subsequentially segmented. Condensates count, area, mean intensity and integrated intensity were measured. Data from the CellProfiler pipeline was further processed by a custom R script for statistics, Mahalanobis distance calculation based on condensates count and integrated intensity z-scores, and for calculation of condensed fractions, as sum of the integrated intensity of the condensates.

### Protein purification

Recombinant mEGFP-β-catenin was expressed in the *E.coli* BL21-DE3 Star strain and grown in Nutrient Broth n1 (Sigma-Aldrich). Following induction with 100 μM IPTG for 14-16 hours at 25°C, cells were harvested by centrifugation at 4000 rpm for 30 min at 4°C, resuspended in lysis buffer (50 mM Tris-HCl pH 7.5, 50 mM NaCl, 5 mM EDTA, 5 mM β-mercaptoethanol and 5% glycerol), lysed by sonication followed centrifugation at 20,000 rpm for 30 min at 4°C. Supernatants were loaded onto Glutathione Sepharose® 4B (Merck) beads in gravity columns (Bio-Rad). Columns were washed twice with 10 volumes of high salt washing buffer (50 mM Tris-HCl pH 7.5, 400 mM NaCl, 5 mM β-mercaptoethanol and 5% glycerol), once with 10 volumes of droplet formation buffer (50 mM Tris-HCl pH 7.5, 125 mM NaCl, 10% glycerol, 1mM DTT) and once with TEV protease buffer (50 mM Tris–HCl pH 8.0, 0.5 mM EDTA and 1 mM DTT). TEV protease His-6 (Protean) cleavage was performed with 1.5 kU of the enzyme in 4 ml of TEV protease buffer overnight, and cleaved proteins were eluted with 2 volumes of TEV protease buffer. Protein was filtered with 50 kDa Amicon Ultra Centrifugal filters (Merck) and dyalised in 50 mM Tris-HCl pH 7.5, 125 mM NaCl, 10% glycerol, 1 mM DTT.

### Droplet assay

Recombinant mEGFP-β-catenin was diluted at 3 μM in droplet formation buffer (50 mM Tris-HCl pH 7.5, 125 mM NaCl, 10% glycerol, 1 mM DTT) together with 10 μM of drug (CYC116 (SelleckChem), CCT129202 (SelleckChem), Pacritinib (SB1518) (SelleckChem), WP1066 (MedChem), YM155 (sepantronium bromide) (SelleckChem), Salermide (MedChem), BIX 01294 (SelleckChem). After addition of 10% PEG-8000 the protein solution was immediately loaded onto microscope slides as a single drop. Slides were imaged with spinning disk confocal microscope (Nikon Ti2) with Apo TIRF 60x Oil DIC N2 lens or spinning disk CSU X1 Nikon Ti with Plan Apo VC 60x N.A. 1.40 oil. Quantifications were performed according to a previously established pipeline, the code for this analysis is available at the following Github link: https://github.com/krishna-shrinivas/2020_Henninger_Oksuz_Shrinivas_RNA_feedback/tree/master/Droplet_analysis^37^. Droplets were segmented by intensity, size and circularity thresholds and their intensity was calculated. The mean intensity of each droplet (C-in) and of the bulk (C-out) were calculated for each channel. The partition ratio was computed as (C-in)/(C-out). The total intensity of each droplet was calculated and then the sum was computed (TC-in) as well as the total bulk intensity of the whole field (TC-out). The condensed fraction was computed as (TC-in)/(TC-out). Quantification data was filtered to exclude droplets with partition ratios lower than 1.05.

### RNA sequencing

4,6 x 10^5^ cells for each well were seeded in 6 well plates. Cells were treated with 1 μM drugs: CYC116 (SelleckChem), CCT129202 (SelleckChem), Pacritinib (SB1518) (SelleckChem), WP1066 (MedChem), YM155 (sepantronium bromide) (SelleckChem), Salermide (MedChem), BIX 01294 (SelleckChem), untreated cells and 0,01% DMSO controls for 12 hours, in triplicate wells. RNA was isolated with RNA isolation kit (QiAgen) and submitted for RNA sequencing and analysis to the USEQ facility. Samples were subjected to stranded polyA library preparation and sequencing with NextSeq2000 50 cycles (P3-1000M clusters). Quality control on the sequence reads from the raw FASTQ files was done with FastQC (v0.11.9). TrimGalore (v0.6.7) as used to trim reads based on quality and adapter presence after which FastQC was again used to check the resulting quality. rRNA reads were filtered out using SortMeRNA (v4.3.6) after which the resulting reads were aligned to the reference genome fasta (GCA_000001405.15_GRCh38_no_alt_plus_hs38d1_analysis_set.fna) using the STAR (v2.7.10b) aligner. Followup QC on the mapped (bam) files was done using Sambamba (v0.8.2), RSeQC (v5.0.1) and PreSeq (v3.2.0). Readcounts were then generated using the Subread FeatureCounts module (v2.0.3) with the Homo_sapiens.GRCh38.106.ncbi.gtf gtf file as annotation, after which normalization was done using the R-package edgeR (v3.40).

Differential gene expression analysis was conducted with a custom script using the DESeq2 R package^38^.

For heatmap visualization data was subjected to log transformation and scaling by row.

### Immunofluorescence

Cells were seeded at 70-80% confluence on 15mm glass cover slides, pre-coated for 5 minutes with 10 μg/mL poly-L-lysine. The following day, cells were treated with 5 µM of the following compounds: CYC116 (SelleckChem), CCT129202 (SelleckChem), Pacritinib (SB1518) (SelleckChem), WP1066 (MedChem), YM155 (sepantronium bromide) (SelleckChem), Salermide (MedChem), and BIX 01294 (SelleckChem) for 1 hour at 37°C. A 0.05% DMSO solution was used as a vehicle control. Subsequently, cells were washed twice with PBS and fixed for 15 minutes at room temperature with 4% paraformaldehyde (PFA) in PBS. After two additional washes with PBS, cells were permeabilized with 0.1% Triton-X100 in PBS for 5 minutes at room temperature. Cover slides were washed twice again with PBS and blocked for 30 minutes with purified 1:10,000 IgG in 0.5% Triton-X100. Cells were then incubated with the appropriate primary antibodies (listed in the table - antibodies) for 1 hour at room temperature or overnight at 4°C. Afterward, cells were washed three times with PBS and incubated with secondary antibodies for 1 hour and 30 minutes at room temperature. Following three more PBS washes, the samples were mounted using slide mounting medium. Images were acquired by scanning a series of Z stacks using a Zeiss LSM880 confocal microscope and analyzed with CellProfiler software.

### Cell viability assay

Cell viability was assessed by propidium iodide exclusion. 1×10^5^ β-catenin KO, mSc-β-catenin^Wt^, mSc-β-catenin^IDRs*^, mSc-β-catenin^nIDR*^, mSc-β-catenin^cIDR*^ cells were seeded for each well of a 12well plate. Cells were treated the day after seeding with DMSO (0,005%, 0,0025%, 0,00125%, 0,00063%, 0,00031%, 0,00016%, 0,00008% and 0,00004%) or YM155 (SelleckChem) (500nM, 250nM, 125nM, 63nM, 31nM, 16nM, 8nM and 4nM) in triplicate. 24 hours after the treatment cells were harvested as follows. Media was collected in glass tubes, cells were washed once with PBS, the PBS was collected in the same previous glass tube. Cells were then trypsinized and collected in the same glass tube. Cells were centrifuged for 5 minutes at 1200 rpm (Beckmann tabletop centrifuge with swingout rotor). Supernatant was removed, and the pelleted cells were resuspended in PBS containing 20 µg/ml propium iodide (PI, Sigma-Aldrich, P4170). At least 5000 events for each sample were measured by flow cytometry with the BD FACSCelesta Cell analyzer (BD biosciences) and analyzed with BD FacsDiva Software

### Organoids viability assay

Organoids were seeded 3-5 days before the experiment in 24well plates. Confluence Organoids were treated with DMSO or YM155 in triplicates for 24 hours and subsequently stained with the two fluorescent dyes 2nM Calcein AM and 5nM Ethidium Homodimer-1 (PTGLab) for 1 hour. Dyes were washed out with PBS and organoids were imaged with the EVOS M5000 microscope (Thermo Fisher Scientific), with a 2X objective. Images were analysed with a custom CellProfiler pipeline in which organoids were segmented and the intensity of the Green (Calcein AM) and Red (Ethidium Homodimer-1) signals were measured. The red/green ratio was calculated, representing the dead/live ratio.

### Dual luciferase assay

For luciferase assays, cells were transfected with a luciferase reporter construct bearing multiple copies of an optimal TCF-binding site (pTOPglow), or the same construct with mutated TCF binding sites (pFOPglow), together with CMV-Renilla. Luciferase activity was analyzed using a luminometer and a dual-luciferase assay kit according to the manufacturer’s protocol (Dual-Luciferase® Reporter Assay System, Promega). Luciferase activity from pTOPglow (TOP) and pFOPglow (FOP) transfected cells was normalized to Renilla activity to control for transfection efficiency. The ratio of the normalized TOP and FOP (TOP/FOP) was then calculated. Calculations and graphs were then made in R.

### Statistical analysis and graphs

All statistical analysis and graphs were made in R (version 4.4.1).

## ACKNOWLEDGMENTS

The authors want to acknowledge Irene Zaalberg for the generation of the APC mutant colon organoid line, Jooske Monster for the generation of the β-catenin KO, mSc-β-catenin^Wt^, mSc-β-catenin^IDRs*^, mSc-β-catenin^nIDR*^ and mSc-β-catenin^cIDR*^ cell lines, Theofilos Chalkiadakis for the help on the RNA-seq data analysis and Marit van Bueren for the help on the image analysis pipeline generation.

