## Supplemental figures for "Condensate screening identifies YM155 as β-catenin condensate inhibitor in colorectal cancer"

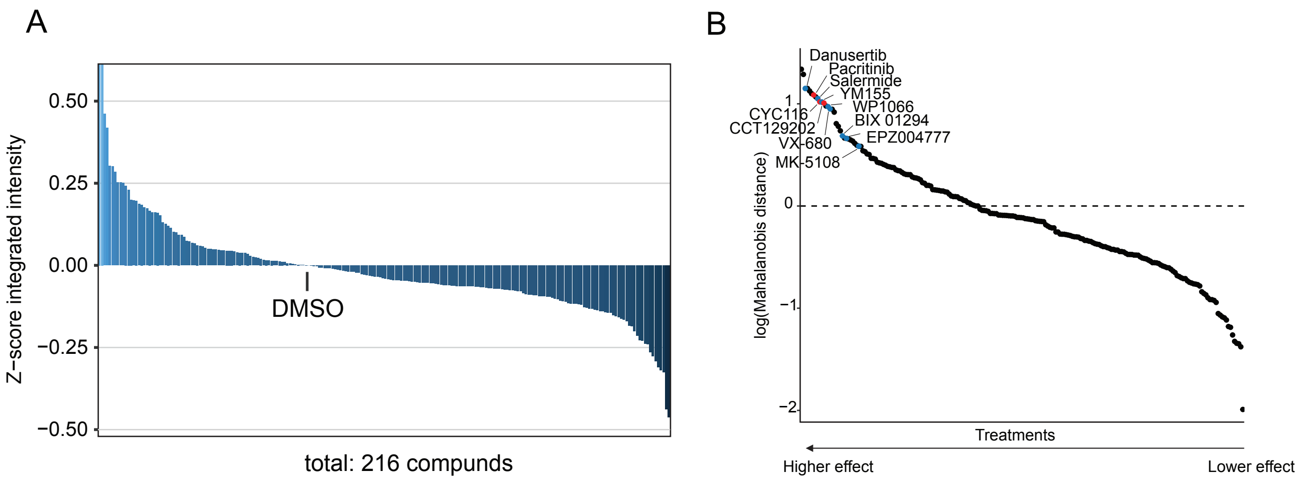

A
0.50
0.25
0.00
−0.25
−0.50
Z−score integrated intensity
DMSO
total: 216 compunds
B
Danusertib
Pacritinib
Salermide
YM155
WP1066
CYC116
CCT129202
BIX 01294
EPZ004777
VX-680
MK-5108
1
0
−1
−2
log(Mahalanobis distance)
Treatments
Higher effect
Lower effect

**Figure S1: Hits identification**

A) Waterfall plot showing the distribution of the 216 compounds and the DMSO control in terms of z-score of the integrated intensity. At least 18 images from 2 replicates were analysed.

B) Mahalanobis distance calculation. On the left side of the plot we can observe the treatments with the highest effect in terms of condensates per nucleus and integrated intensity combined. Reducers hits compounds are indicated in blue, while enhancers are indicated in red. At least 18 images from 2 replicates were analysed.

A

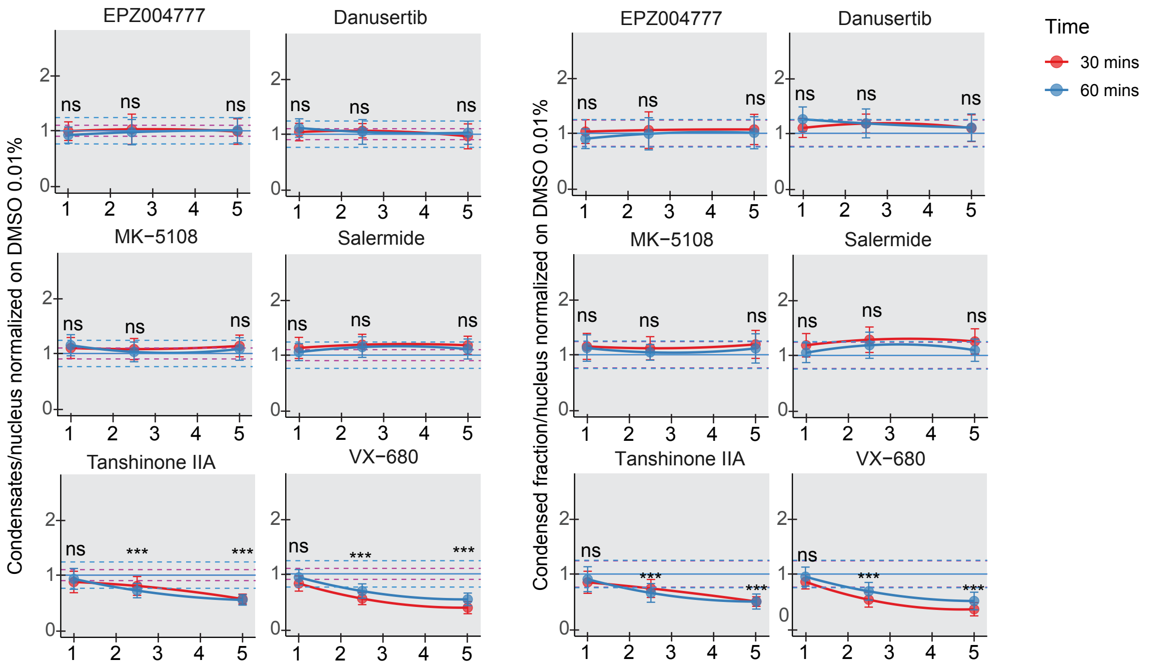
EPZ004777
Danusertib
MK−5108
Salermide
Tanshinone IIA
VX−680
Condensates/nucleus normalized on DMSO 0.01%
Condensed fraction/nucleus normalized on DMSO 0.01%
ns
***
Time
30 mins
60 mins

B

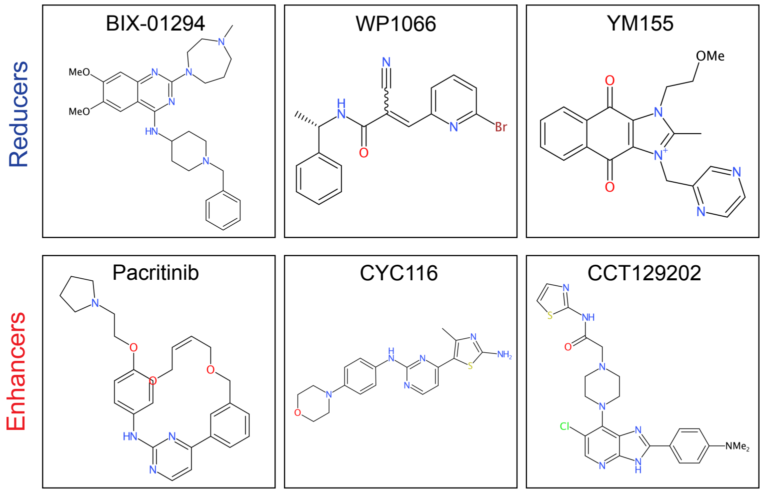
Reducers
Enhancers
BIX-01294
WP1066
YM155
Pacritinib
CYC116
CCT129202

**Figure S2: Hits validation and specificity assessment**

A) Left: Quantification of the average number of β-catenin condensates per nucleus of HCT 116 NECO cells after 30 minutes (red) and 1 hour (blue) treatment with 0,01% DMSO and EPZ004777, Danusertib, MK-5108, Salermide and VX-680 (1 μM, 2.5 μM and 5 μM). Tanshinone IIA (1 μM, 2.5 μM and 5 μM) was used as positive control.
Right: Quantification of the average condensed fraction of β-catenin condensates per nucleus in HCT 116 NECO cells after 30 minutes (red) and 1 hour (blue) treatment with 0,01% DMSO and EPZ004777, Danusertib, MK-5108, Salermide and VX-680 (1 μM, 2.5 μM and 5 μM). Tanshinone IIA (1 μM, 2.5 μM and 5 μM) was used as positive control.
Continuous lines at y=1 represent DMSO controls, while dotted red and blue lines represent the error bars interval for DMSO at 30 minutes and 1 hour, respectively.
At least five biological replicates for each treatment for each concentration were considered. 9 images were acquired for each biological replicate. ANOVA and Tukey's HSD multiple comparison test were performed for each concentration. ns: not significative, ***$p < 0.001$.

B) Chemical structures of the six selected compounds obtained with rcdk package in R.[19]

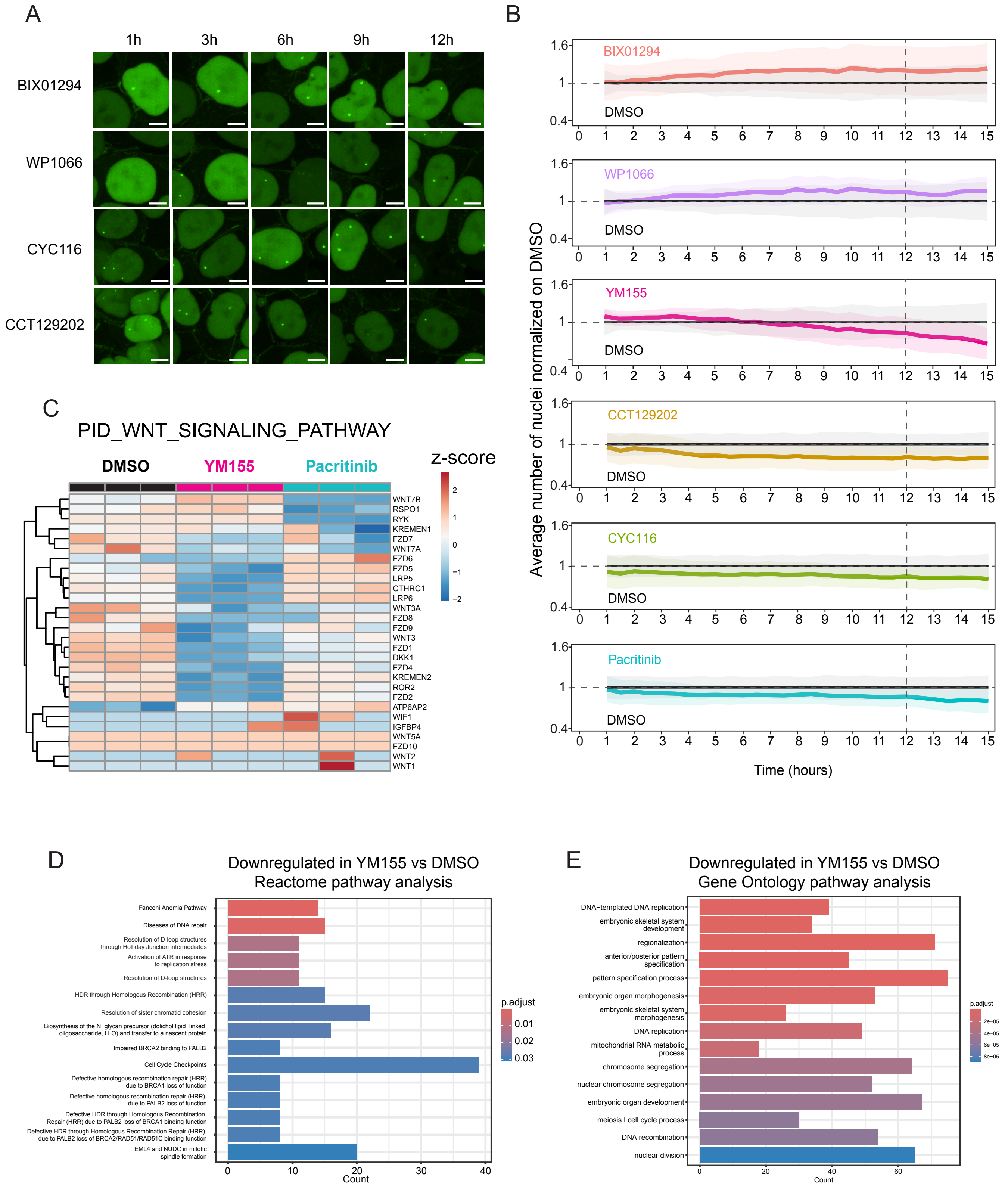
A
1h
3h
6h
9h
12h
BIX01294
WP1066
CYC116
CCT129202
B
Average number of nuclei normalized on DMSO
BIX01294
WP1066
YM155
CCT129202
CYC116
Pacritinib
DMSO
1.6
1
0.4
0 1 2 3 4 5 6 7 8 9 10 11 12 13 14 15
Time (hours)
C
PID_WNT_SIGNALING_PATHWAY
DMSO
YM155
Pacritinib
z-score
2
1
0
−1
−2
WNT7B
RSPO1
RYK
KREMEN1
FZD7
WNT7A
FZD6
FZD5
LRP5
CTHRC1
LRP6
WNT3A
FZD8
FZD9
WNT3
FZD1
DKK1
FZD4
KREMEN2
ROR2
FZD2
ATP6AP2
WIF1
IGFBP4
WNT5A
FZD10
WNT2
WNT1
D
Downregulated in YM155 vs DMSO
Reactome pathway analysis
Fanconi Anemia Pathway
Diseases of DNA repair
Resolution of D-loop structures through Holliday Junction intermediates
Activation of ATR in response to replication stress
Resolution of D-loop structures
HDR through Homologous Recombination (HRR)
Resolution of sister chromatid cohesion
Biosynthesis of the N−glycan precursor (dolichol lipid−linked oligosaccharide, LLO) and transfer to a nascent protein
Impaired BRCA2 binding to PALB2
Cell Cycle Checkpoints
Defective homologous recombination repair (HRR) due to BRCA1 loss of function
Defective homologous recombination repair (HRR) due to PALB2 loss of function
Defective HDR through Homologous Recombination Repair (HRR) due to PALB2 loss of BRCA1 binding function
Defective HDR through Homologous Recombination Repair (HRR) due to PALB2 loss of BRCA2/RAD51/RAD51C binding function
EML4 and NUDC in mitotic spindle formation
0 10 20 30 40
Count
p.adjust
0.01
0.02
0.03
E
Downregulated in YM155 vs DMSO
Gene Ontology pathway analysis
DNA−templated DNA replication
embryonic skeletal system development
regionalization
anterior/posterior pattern specification
pattern specification process
embryonic organ morphogenesis
embryonic skeletal system morphogenesis
DNA replication
mitochondrial RNA metabolic process
chromosome segregation
nuclear chromosome segregation
embryonic organ development
meiosis I cell cycle process
DNA recombination
nuclear division
0 20 40 60
Count
p.adjust
2e−05
4e−05
6e−05
8e−05

**Figure S4: Transcriptome analysis and live imaging upon treatments with selected compounds**

A) Representative images of live HCT 116 NECO cells at 1 hour, 3 hours, 6 hours, 9 hours and 12 hours timepoints after 1µM BIX01294, 1µM WP1066, 1µM CYC116 and 1µM CCT129202 treatments. Scalebar 5µm.
B) Quantification of the average number of nuclei per image over time upon the treatment with 0,01% DMSO and six compounds: YM155, WP1066, BIX-01294, Pacritinib, CYC116 and CCT129202 (1µM). A total of 16 images for treatment were collected, from 4 biological replicates from two independent experiments.
C) Heatmap showing the relative gene expression changes (z-scores) in the gene involved in the PID_WNT_SIGNALING_PATHWAY from the Panther database in the samples treated with 0,01% DMSO, 1µM YM155 or 1µM Pacritinib for 12 hours.
D) Enriched pathways from Reactome database pathways analysis on the significatively downregulated genes upon 12 hours 1µM YM155 treatment. Treatments were performed in triplicates.
E) Enriched pathways from Gene Ontology database pathways analysis on the significatively downregulated genes upon 12 hours 1µM YM155 treatment. Treatments were performed in triplicates.

A

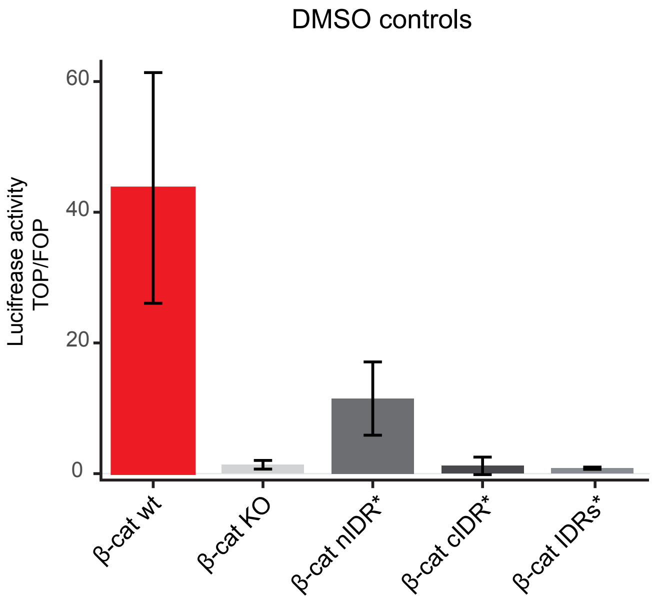
DMSO controls
Luciferase activity TOP/FOP
60
40
20
0
β-cat wt
β-cat KO
β-cat nIDR*
β-cat cIDR*
β-cat IDRs*

**Figure S5: Basal WNT-transcription of the HCT 116 cell lines**

A) Luciferase assay experiment showing the basal level of transctiption for the 0,00004% DMSO controls for the cell lines mSc-β-catenin$^{Wt}$ (β-cat$^{Wt}$), mSc-β-catenin$^{nIDR*}$ (β-cat$^{nIDR*}$), mSc-β-catenin$^{IDRs*}$ (β-cat$^{IDRs*}$), β-catenin KO (β-cat KO), mSc-β-catenin$^{cIDR*}$ (β-cat$^{cIDR*}$). Six replicates from 2 independent experiments were considered.
