## Supplemental Table 1 for "Condensate screening identifies YM155 as β-catenin condensate inhibitor in colorectal cancer"

**Supplementary Table 1: rcdk molecular descriptors**

**Descriptor 2**

| compound | Pacritinib | CYC116 | CCT129202 | WP1066 | BIX01294 | YM155 |
|---|---|---|---|---|---|---|
| BCUTw.1l | 11.6867788 | 11.6995372 | 11.6939429 | 11.6939331 | 11.6828701 | 11.6852286 |
| BCUTw.1h | 16.0021655 | 31.9743259 | 34.9693814 | 78.9185182 | 16.0016758 | 16.0074277 |
| BCUTc.1l | -0.4180097 | -0.4225159 | -0.3895926 | -0.4272763 | -0.4080948 | -0.447366 |
| BCUTc.1h | 0.35499103 | 0.37006331 | 0.41943405 | 0.42142945 | 0.40606842 | 0.42442493 |
| BCUTp.1l | 6.61362985 | 5.72851452 | 5.16378454 | 4.68558986 | 5.02311745 | 4.55801884 |
| BCUTp.1h | 10.1533862 | 11.9030928 | 11.7441845 | 9.97156642 | 11.0583561 | 11.6259962 |

**Descriptor 3**

| compound | Pacritinib | CYC116 | CCT129202 | WP1066 | BIX01294 | YM155 |
|---|---|---|---|---|---|---|
| Fsp3 | 0.35714286 | 0.27777778 | 0.30434783 | 0.11764706 | 0.5 | 0.25 |
| XLogP | 4.157 | 1.557 | 2.859 | 3.3 | 4.184 | 0.144 |
| MW | 472.579737 | 368.45777 | 497.017498 | 356.216723 | 490.641383 | 363.39062 |
| LipinskiFailur | 0 | 0 | 0 | 0 | 0 | 0 |
| nRotB | 4 | 4 | 7 | 5 | 7 | 5 |
| MLogP | 3.77 | 2.56 | 2.78 | 2.78 | 3.66 | 2.89 |
| nAtomLAC | 2 | 0 | 2 | 3 | 0 | 2 |
| nAtomP | 20 | 20 | 17 | 13 | 14 | 15 |
| nAtomLC | 3 | 0 | 3 | 7 | 2 | 4 |
| nB | 39 | 29 | 38 | 23 | 40 | 30 |
| nBase | 1 | 0 | 1 | 0 | 2 | 1 |
| nAtom | 67 | 46 | 59 | 36 | 74 | 46 |
| nAromBond | 18 | 17 | 21 | 12 | 17 | 0 |
| naAromAtom | 18 | 17 | 20 | 12 | 16 | 0 |
| ALogP | 4.547 | 2.7438 | 2.9725 | 3.5665 | 4.1004 | -2.7177 |
| ALogp2 | 20.675209 | 7.52843844 | 8.83575625 | 12.7199223 | 16.8132802 | 7.38589329 |
| AMR | 139.425 | 105.2172 | 136.447 | 89.7993 | 147.588 | 104.0131 |

**Descriptor 4**

| compound | Pacritinib | CYC116 | CCT129202 | WP1066 | BIX01294 | YM155 |
|---|---|---|---|---|---|---|
| nSmallRings | 4 | 4 | 5 | 2 | 5 | 4 |
| nAromRings | 3 | 3 | 4 | 2 | 3 | 4 |
| nRingBlocks | 2 | 4 | 4 | 2 | 4 | 2 |
| nAromBlocks | 3 | 3 | 3 | 2 | 2 | 2 |
| nRings3 | 0 | 0 | 0 | 0 | 0 | 0 |
| nRings4 | 0 | 0 | 0 | 0 | 0 | 0 |
| nRings5 | 1 | 1 | 2 | 0 | 0 | 1 |
| nRings6 | 3 | 3 | 3 | 2 | 4 | 3 |
| nRings7 | 0 | 0 | 0 | 0 | 1 | 0 |
| nRings8 | 0 | 0 | 0 | 0 | 0 | 0 |
| nRings9 | 0 | 0 | 0 | 0 | 0 | 0 |
| tpsaEfficienc | 0.14555929 | 0.31898024 | 0.24492294 | 0.18527906 | 0.13458952 | 0.20471121 |
| Zagreb | 180 | 138 | 184 | 106 | 190 | 146 |
| WPATH | 3973 | 1895 | 3956 | 1159 | 4277 | 1672 |
| WPOL | 51 | 38 | 53 | 30 | 61 | 49 |
| WTPT.1 | 73.0411982 | 53.8107775 | 70.2240947 | 43.9248634 | 74.6957992 | 55.6486117 |
| WTPT.2 | 2.08689138 | 2.06964529 | 2.06541455 | 1.9965847 | 2.07488331 | 2.06105969 |
| WTPT.3 | 22.0204898 | 24.4011164 | 33.6817728 | 13.5417386 | 25.4864778 | 21.0884133 |
| WTPT.4 | 9.21107929 | 3.00841145 | 2.54661093 | 2.53578035 | 5.62692553 | 7.9695414 |
| WTPT.5 | 12.8094105 | 18.3581029 | 25.5482189 | 8.48247181 | 19.8595523 | 13.1188719 |
| VAdjMat | 6.28540222 | 5.857981 | 6.24792751 | 5.52356196 | 6.32192809 | 5.9068906 |
| VABC | | | | | | |
| TopoPSA | 68.74 | 117.43 | 121.52 | 65.78 | 65.99 | 74.34 |
| topoShape | 0.88888889 | 0.875 | 0.9 | 1 | 0.88888889 | 0.83333333 |
| geomShape | #NUM! | #NUM! | #NUM! | #NUM! | #NUM! | #NUM! |
| PetitjeanNur | 0.47058824 | 0.46666667 | 0.47368421 | 0.5 | 0.47058824 | 0.45454545 |
| MDEC.11 | 0 | 0 | 0.5 | 0 | 0.34456668 | 0.16666667 |
| MDEC.12 | 0 | 1.14763975 | 2.8506842 | 2.01478958 | 5.75735205 | 3.33623015 |
| MDEC.13 | 0 | 2.00601929 | 2.35766486 | 1.85314401 | 4.04970255 | 3.47583112 |
| MDEC.14 | 0 | 0 | 0 | 0 | 0 | 0 |
| MDEC.22 | 0 | 11.4048912 | 11.5589319 | 10.0251357 | 23.6182888 | 9.32887171 |
| MDEC.23 | 28.5614146 | 14.1115582 | 21.6247518 | 15.4713816 | 25.2719641 | 20.6666744 |
| MDEC.24 | 0 | 0 | 0 | 0 | 0 | 0 |
| MDEC.33 | 5.56777401 | 5.91002885 | 8.13799647 | 4.77054201 | 8.04006764 | 11.9713474 |
| MDEC.34 | 0 | 0 | 0 | 0 | 0 | 0 |
| MDEC.44 | 0 | 0 | 0 | 0 | 0 | 0 |
| MDEO.11 | 0 | 0 | 0 | 0 | 0 | 0.2 |
| MDEO.12 | 0 | 0 | 0 | 0 | 0 | 0.3086067 |
| MDEO.22 | 0.53132928 | 0 | 0 | 0 | 0.33333333 | 0 |
| MDEN.11 | 0 | 0 | 0 | 0 | 0 | 0 |
| MDEN.12 | 0 | 0.85018017 | 0 | 0.4472136 | 0 | 0 |
| MDEN.13 | 0 | 0.08333333 | 0 | 0 | 0 | 0 |
| MDEN.22 | 1.5 | 1.85314401 | 1.76432711 | 0.2 | 1.19055079 | 0.33333333 |
| MDEN.23 | 0.3231652 | 0.55516273 | 2.4241957 | 0 | 2.07742242 | 0.91829954 |
| MDEN.33 | 0 | 0 | 0.43679023 | 0 | 0.46764893 | 0.5 |
| khs.sLi | 0 | 0 | 0 | 0 | 0 | 0 |

| | | | | | | |
|---|---|---|---|---|---|---|
| khs.ssBe | 0 | 0 | 0 | 0 | 0 | 0 |
| khs.ssssBe | 0 | 0 | 0 | 0 | 0 | 0 |
| khs.ssBH | 0 | 0 | 0 | 0 | 0 | 0 |
| khs.sssB | 0 | 0 | 0 | 0 | 0 | 0 |
| khs.ssssB | 0 | 0 | 0 | 0 | 0 | 0 |
| khs.sCH3 | 0 | 1 | 2 | 1 | 3 | 2 |
| khs.dCH2 | 0 | 0 | 0 | 0 | 0 | 0 |
| khs.ssCH2 | 10 | 4 | 5 | 0 | 10 | 3 |
| khs.tCH | 0 | 0 | 0 | 0 | 0 | 0 |
| khs.dsCH | 2 | 0 | 0 | 1 | 0 | 0 |
| khs.aaCH | 9 | 6 | 7 | 8 | 7 | 7 |
| khs.sssCH | 0 | 0 | 0 | 1 | 1 | 0 |
| khs.ddC | 0 | 0 | 0 | 0 | 0 | 0 |
| khs.tsC | 0 | 0 | 0 | 1 | 0 | 0 |
| khs.dssC | 0 | 0 | 1 | 2 | 0 | 2 |
| khs.aasC | 7 | 7 | 6 | 3 | 5 | 6 |
| khs.aaaC | 0 | 0 | 2 | 0 | 2 | 0 |
| khs.ssssC | 0 | 0 | 0 | 0 | 0 | 0 |
| khs.sNH3 | 0 | 0 | 0 | 0 | 0 | 0 |
| khs.sNH2 | 0 | 1 | 0 | 0 | 0 | 0 |
| khs.ssNH2 | 0 | 0 | 0 | 0 | 0 | 0 |
| khs.dNH | 0 | 0 | 0 | 0 | 0 | 0 |
| khs.ssNH | 1 | 1 | 1 | 1 | 1 | 0 |
| khs.aaNH | 0 | 0 | 1 | 0 | 0 | 0 |
| khs.tN | 0 | 0 | 0 | 1 | 0 | 0 |
| khs.sssNH | 0 | 0 | 0 | 0 | 0 | 0 |
| khs.dsN | 0 | 0 | 0 | 0 | 0 | 0 |
| khs.aaN | 2 | 3 | 3 | 1 | 2 | 2 |
| khs.sssN | 1 | 1 | 3 | 0 | 3 | 0 |
| khs.ddsN | 0 | 0 | 0 | 0 | 0 | 0 |
| khs.aasN | 0 | 0 | 0 | 0 | 0 | 2 |
| khs.ssssN | 0 | 0 | 0 | 0 | 0 | 0 |
| khs.sOH | 0 | 0 | 0 | 0 | 0 | 0 |
| khs.dO | 0 | 0 | 1 | 1 | 0 | 2 |
| khs.ssO | 3 | 1 | 0 | 0 | 2 | 1 |
| khs.aaO | 0 | 0 | 0 | 0 | 0 | 0 |
| khs.sF | 0 | 0 | 0 | 0 | 0 | 0 |
| khs.sSiH3 | 0 | 0 | 0 | 0 | 0 | 0 |
| khs.ssSiH2 | 0 | 0 | 0 | 0 | 0 | 0 |
| khs.sssSiH | 0 | 0 | 0 | 0 | 0 | 0 |
| khs.ssssSi | 0 | 0 | 0 | 0 | 0 | 0 |
| khs.sPH2 | 0 | 0 | 0 | 0 | 0 | 0 |
| khs.ssPH | 0 | 0 | 0 | 0 | 0 | 0 |
| khs.sssP | 0 | 0 | 0 | 0 | 0 | 0 |
| khs.dsssP | 0 | 0 | 0 | 0 | 0 | 0 |
| khs.sssssP | 0 | 0 | 0 | 0 | 0 | 0 |
| khs.sSH | 0 | 0 | 0 | 0 | 0 | 0 |
| khs.dS | 0 | 0 | 0 | 0 | 0 | 0 |

| | | | | | | |
|---|---|---|---|---|---|---|
| khs.ssS | 0 | 0 | 0 | 0 | 0 | 0 |
| khs.aaS | 0 | 1 | 1 | 0 | 0 | 0 |
| khs.dssS | 0 | 0 | 0 | 0 | 0 | 0 |
| khs.ddssS | 0 | 0 | 0 | 0 | 0 | 0 |
| khs.sCl | 0 | 0 | 1 | 0 | 0 | 0 |
| khs.sGeH3 | 0 | 0 | 0 | 0 | 0 | 0 |
| khs.ssGeH2 | 0 | 0 | 0 | 0 | 0 | 0 |
| khs.sssGeH | 0 | 0 | 0 | 0 | 0 | 0 |
| khs.ssssGe | 0 | 0 | 0 | 0 | 0 | 0 |
| khs.sAsH2 | 0 | 0 | 0 | 0 | 0 | 0 |
| khs.ssAsH | 0 | 0 | 0 | 0 | 0 | 0 |
| khs.sssAs | 0 | 0 | 0 | 0 | 0 | 0 |
| khs.sssdAs | 0 | 0 | 0 | 0 | 0 | 0 |
| khs.sssssAs | 0 | 0 | 0 | 0 | 0 | 0 |
| khs.sSeH | 0 | 0 | 0 | 0 | 0 | 0 |
| khs.dSe | 0 | 0 | 0 | 0 | 0 | 0 |
| khs.ssSe | 0 | 0 | 0 | 0 | 0 | 0 |
| khs.aaSe | 0 | 0 | 0 | 0 | 0 | 0 |
| khs.dssSe | 0 | 0 | 0 | 0 | 0 | 0 |
| khs.ddssSe | 0 | 0 | 0 | 0 | 0 | 0 |
| khs.sBr | 0 | 0 | 0 | 1 | 0 | 0 |
| khs.sSnH3 | 0 | 0 | 0 | 0 | 0 | 0 |
| khs.ssSnH2 | 0 | 0 | 0 | 0 | 0 | 0 |
| khs.sssSnH | 0 | 0 | 0 | 0 | 0 | 0 |
| khs.ssssSn | 0 | 0 | 0 | 0 | 0 | 0 |
| khs.sI | 0 | 0 | 0 | 0 | 0 | 0 |
| khs.sPbH3 | 0 | 0 | 0 | 0 | 0 | 0 |
| khs.ssPbH2 | 0 | 0 | 0 | 0 | 0 | 0 |
| khs.sssPbH | 0 | 0 | 0 | 0 | 0 | 0 |
| khs.ssssPb | 0 | 0 | 0 | 0 | 0 | 0 |
| Kier1 | 26.6009204 | 19.3222354 | 25.6412742 | 18.3402647 | 27.5625 | 20.28 |
| Kier2 | 14.2352941 | 9 | 11.5884774 | 9.33333333 | 13.3752066 | 8.78853434 |
| Kier3 | 8.24047337 | 4.89940828 | 6.12345679 | 5.86419753 | 7.15856634 | 3.77324263 |
| HybRatio | 0.35714286 | 0.27777778 | 0.30434783 | 0.125 | 0.5 | 0.25 |
| fragC | 3851.07 | 1751.08 | 2847.11 | 907.05 | 4824.08 | 1699.07 |
| FMF | 1 | 0.92307692 | 0.85294118 | 0.77272727 | 0.86111111 | 0.74074074 |
| ECCEN | 1003 | 669 | 1073 | 427 | 1046 | 497 |
| SP.0 | 23.7106852 | 17.9325107 | 23.6561253 | 15.949383 | 24.9072021 | 18.9658908 |
| SP.1 | 17.3316325 | 12.6866732 | 16.4743669 | 10.5965555 | 17.6228561 | 13.130229 |
| SP.2 | 14.8004182 | 11.4250237 | 15.2141439 | 9.13019337 | 15.4592555 | 11.5538929 |
| SP.3 | 12.3278485 | 9.48303399 | 12.7380297 | 7.20833395 | 13.2804133 | 10.4449625 |
| SP.4 | 10.3551157 | 8.16751239 | 10.868005 | 5.67783553 | 11.0560992 | 9.20969471 |
| SP.5 | 8.10198404 | 6.40940909 | 8.32059004 | 4.06556925 | 9.50931911 | 8.05546845 |
| SP.6 | 5.7529239 | 3.89787513 | 6.11909491 | 2.3328329 | 6.992441 | 5.65118825 |
| SP.7 | 4.50746522 | 2.96010375 | 4.87996057 | 1.48929676 | 5.26398783 | 4.24620598 |
| VP.0 | 19.9883064 | 15.2917266 | 20.5271539 | 13.5401392 | 21.7424345 | 15.4362647 |
| VP.1 | 12.3135486 | 9.26644116 | 12.2210837 | 7.49367639 | 13.0276961 | 9.12683545 |
| VP.2 | 8.81223811 | 7.0340689 | 9.62906404 | 5.47330282 | 9.92078927 | 7.03423023 |

| | | | | | | |
|---|---|---|---|---|---|---|
| VP.3 | 6.17983069 | 5.21013132 | 6.79890235 | 3.48277047 | 7.36902342 | 5.60824425 |
| VP.4 | 4.34880418 | 3.7098678 | 4.85259251 | 2.25763915 | 5.21173247 | 4.27866939 |
| VP.5 | 2.65326488 | 2.31814996 | 3.08609275 | 1.33715199 | 3.86467153 | 3.20109843 |
| VP.6 | 1.54126613 | 1.08494518 | 1.87116361 | 0.67927033 | 2.39383919 | 2.07819388 |
| VP.7 | 0.96251435 | 0.69149018 | 1.27657021 | 0.3264655 | 1.47669317 | 1.37014809 |
| SPC.4 | 3.42035845 | 3.43301833 | 5.29920291 | 2.7572924 | 4.80109203 | 4.62200141 |
| SPC.5 | 5.48694722 | 5.65541127 | 8.37976019 | 3.9695585 | 7.56124416 | 8.55563538 |
| SPC.6 | 7.11882449 | 7.24989415 | 11.0758388 | 4.88556449 | 10.8115081 | 14.1447737 |
| VPC.4 | 1.38470546 | 1.73812914 | 2.35938185 | 1.10335854 | 2.1759122 | 2.22436909 |
| VPC.5 | 1.84993264 | 2.36304689 | 3.07073443 | 1.29274249 | 2.80693485 | 3.54057701 |
| VPC.6 | 1.77571155 | 2.37191248 | 3.37095789 | 1.2875542 | 3.3069707 | 4.90095771 |
| SC.3 | 1.48316325 | 1.56870845 | 2.34648623 | 1.29753713 | 2.03713033 | 1.54802769 |
| SC.4 | 0 | 0 | 0 | 0 | 0 | 0 |
| SC.5 | 0.16666667 | 0.24759976 | 0.49752528 | 0.23570226 | 0.30274943 | 0.62588418 |
| SC.6 | 0 | 0 | 0 | 0 | 0 | 0 |
| VC.3 | 0.71816904 | 0.80723244 | 1.24776265 | 0.69184431 | 1.11090547 | 0.8441476 |
| VC.4 | 0 | 0 | 0 | 0 | 0 | 0 |
| VC.5 | 0.04557284 | 0.14526159 | 0.1827166 | 0.06284391 | 0.0710209 | 0.26233107 |
| VC.6 | 0 | 0 | 0 | 0 | 0 | 0 |
| SCH.3 | 0 | 0 | 0 | 0 | 0 | 0 |
| SCH.4 | 0 | 0 | 0 | 0 | 0 | 0 |
| SCH.5 | 0.14433757 | 0.09622504 | 0.24056261 | 0 | 0 | 0.06415003 |
| SCH.6 | 0.33677012 | 0.51673438 | 0.50343679 | 0.18539541 | 0.29650652 | 0.45137839 |
| SCH.7 | 0.43058253 | 0.55890532 | 0.7298591 | 0.20118446 | 0.4898722 | 0.83572862 |
| VCH.3 | 0 | 0 | 0 | 0 | 0 | 0 |
| VCH.4 | 0 | 0 | 0 | 0 | 0 | 0 |
| VCH.5 | 0.1118034 | 0.06846532 | 0.11923794 | 0 | 0 | 0.03952847 |
| VCH.6 | 0.14755765 | 0.23231444 | 0.1800096 | 0.05359159 | 0.12995807 | 0.19760643 |
| VCH.7 | 0.14495851 | 0.19439884 | 0.20636893 | 0.07319883 | 0.16361676 | 0.34084181 |
| C1SP1 | 0 | 0 | 0 | 1 | 0 | 0 |
| C2SP1 | 0 | 0 | 0 | 0 | 0 | 0 |
| C1SP2 | 1 | 1 | 6 | 2 | 1 | 4 |
| C2SP2 | 13 | 10 | 8 | 10 | 10 | 9 |
| C3SP2 | 3 | 0 | 1 | 2 | 2 | 2 |
| C1SP3 | 8 | 5 | 5 | 1 | 7 | 4 |
| C2SP3 | 2 | 0 | 0 | 1 | 4 | 0 |
| C3SP3 | 0 | 0 | 0 | 0 | 0 | 0 |
| C4SP3 | 0 | 0 | 0 | 0 | 0 | 0 |
| ATSp1 | 2348.15502 | 1908.3495 | 2620.77353 | 1441.94561 | 2622.304 | 1956.89228 |
| ATSp2 | 2689.3189 | 2231.01258 | 3096.89431 | 1613.66536 | 3075.95243 | 2394.19577 |
| ATSp3 | 3548.07409 | 3062.5272 | 4368.65867 | 2061.26297 | 4286.55741 | 3597.94458 |
| ATSp4 | 3572.98037 | 2846.94848 | 4258.09886 | 1994.9904 | 4725.06332 | 3949.52668 |
| ATSp5 | 3488.82975 | 2444.99888 | 4028.49038 | 1714.12639 | 4182.19927 | 3702.33195 |
| ATSm1 | 38.7633283 | 35.0621039 | 51.4950144 | 67.1125007 | 39.7088199 | 30.7633283 |
| ATSm2 | 42.4881893 | 34.9979376 | 46.7808898 | 29.8156771 | 43.8210969 | 32.9901872 |
| ATSm3 | 55.7340402 | 51.4985874 | 68.7570063 | 44.4597325 | 59.9286542 | 47.1808053 |
| ATSm4 | 56.2041087 | 46.3330343 | 66.4905447 | 43.6310823 | 67.1188198 | 54.7023527 |
| ATSm5 | 56.75748 | 40.3581898 | 69.2864972 | 41.2121604 | 60.1189553 | 56.1138266 |

| | | | | | | |
|---|---|---|---|---|---|---|
| ATSc1 | 0.77714997 | 0.54979846 | 0.74881067 | 0.37645313 | 0.76279541 | 0.81716507 |
| ATSc2 | -0.5145717 | -0.373673 | -0.4545747 | -0.1610338 | -0.5224841 | -0.3773802 |
| ATSc3 | 0.14046958 | 0.05432678 | -0.0928883 | 0.05183721 | 0.18987968 | -0.2592377 |
| ATSc4 | 0.01521487 | 0.06855495 | 0.24967856 | -0.1566903 | -0.0739646 | 0.31459686 |
| ATSc5 | -0.0836091 | -0.0460728 | 0.01932102 | -0.0037998 | -0.0575563 | -0.0399153 |

**Descriptor 5**

| compound | Pacritinib | CYC116 | CCT129202 | WP1066 | BIX01294 | YM155 |
|---|---|---|---|---|---|---|
| tpsaEfficienc | 0.14555929 | 0.31898024 | 0.24492294 | 0.18527906 | 0.13458952 | 0.20471121 |
| TopoPSA | 68.74 | 117.43 | 121.52 | 65.78 | 65.99 | 74.34 |
| nHBDon | 1 | 2 | 2 | 1 | 1 | 0 |
| nHBAcc | 6 | 7 | 8 | 4 | 6 | 6 |
| bpol | 46.010624 | 32.00014 | 42.208175 | 20.192898 | 54.613866 | 31.202933 |
| apol | 77.423376 | 55.31786 | 71.831825 | 46.407102 | 82.822134 | 54.675067 |

**Descriptor 6**

| compound | Pacritinib | CYC116 | CCT129202 | WP1066 | BIX01294 | YM155 |
|---|---|---|---|---|---|---|
| topoShape | 0.88888889 | 0.875 | 0.9 | 1 | 0.88888889 | 0.83333333 |
